# A Phylogenetic Approach to Inferring the Order in Which Mutations Arise during Cancer Progression

**DOI:** 10.1101/2020.05.06.081398

**Authors:** Yuan Gao, Jeff Gaither, Julia Chifman, Laura Kubatko

## Abstract

Although the role of evolutionary process in cancer progression is widely accepted, increasing attention is being given to the evolutionary mechanisms that can lead to differences in clinical outcome. Recent studies suggest that the temporal order in which somatic mutations accumulate during cancer progression is important. Single-cell sequencing provides a unique opportunity to examine the mutation order during cancer progression. However, the errors associated with single-cell sequencing complicate this task. We propose a new method for inferring the order in which somatic mutations arise within a tumor using noisy single-cell sequencing data that incorporates the errors that arise from the data collection process. Using simulation, we show that our method outperforms existing methods for identifying mutation order in most cases, especially when the number of cells is large. Our method also provides a means to quantify the uncertainty in the inferred mutation order along a fixed phylogeny. We apply our method to empirical data from colorectal and prostate cancer patients.

## 1. Introduction

Cancer progression is a dynamic evolutionary process that occurs among the individual cells within each patient’s tumor. Cancer develops from a single cell in normal tissue whose genetic alterations endow a growth advantage over the surrounding cells, allowing that cell to replicate and to expand, resulting in the formation of a clonal population of identical cells. Cells within this clonal population may then undergo their own somatic mutations, followed by replication and formation of subclones. During this complex process, many competitive and genetically diverse subpopulations may be formed, resulting in intratumoral heterogeneity (ITH) depicted in Fig. 1(a) (O’Sullivan *and others*, 2003; Ishwaran *and others*, 2009; Jamal-Hanjani *and others*, 2017; Ascolani and Liò, 2019). Ortmann *and others* (2015) demonstrate that the type of malignancy and the response to treatment of myeloproliferative neoplasm patients are affected by the order in which somatic mutations arose within the patients’ tumors. Though this study is specific to one type of cancer, the timing and organization of somatic mutations are crucial to clinical outcomes for other cancers as well. Determining the temporal order of mutations required for tumor progression is thus critical, especially in the field of targeted therapy. However, this information cannot be observed directly, since genomic data are most often collected at one snapshot in time. Consequently, use of computational methods that infer the order of mutations from DNA sequence data is the approach of choice.

**Fig. 1:**
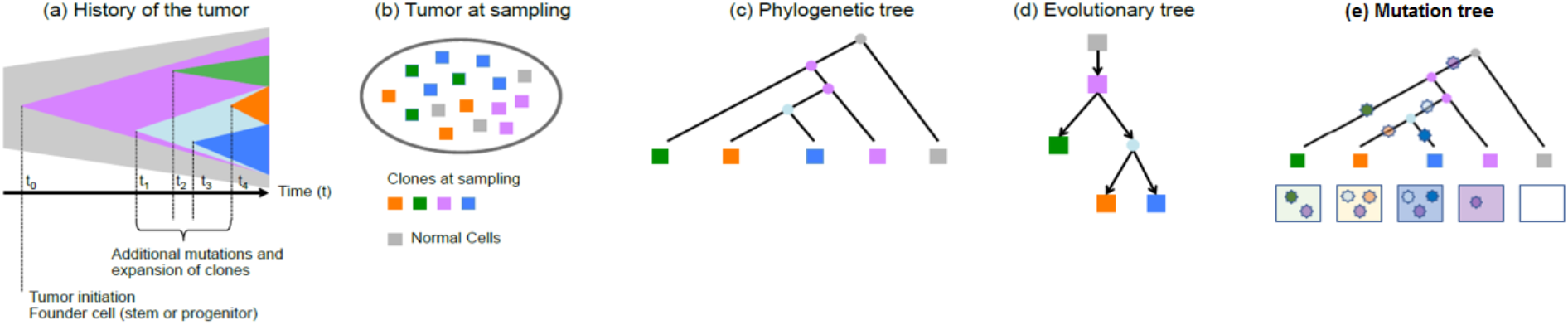
Pictorial representation of tumor evolution. (a) - (b) A pictorial representation of the evolution of a tumor from the first initiating mutation to the heterogeneous tissue at the time of sampling, which consists of four different clones and normal tissue. (c) A phylogenetic tree with single cells as the tips. (d) A clonal lineage tree inferred from sampled cells where each node represents a subclone (cluster of cells). (e) A mutation tree inferred from sampled cells where each star represents the occurrence of one mutation. The box underneath each tip shows which mutations are present in the cell represented by the tip.

Most studies on cancer phylogenetics utilize bulk high-throughput sequencing data, but signals from bulk sequencing only reflect the overall characteristics of a population of sequenced cells, rather than the characteristics of individual cells. Variation in the mutational signatures among different cells in a tumor is thus difficult to evaluate from bulk sequencing data. Single-cell sequencing (SCS) is promising because it enables sequencing of individual cells, thus providing the highest possible resolution available on the mutational history of cancer. However, the high error probabilities associated with SCS data complicate the development of methods for inference of the mutational history. The whole-genome amplification (WGA) process used to produce SCS data results in a variety of errors, including allelic dropout (ADO) errors, false positives (FPs), non-uniform coverage distribution, and low coverage regions. ADO contributes a considerable number of false negatives (FNs) to point mutations (Navin, 2014).

Recently, several studies have proposed various mathematical methods to infer mutation order (Fig. 1(c) - Fig. 1(e)) from data arising from single-cell somatic mutations. Of particular interest are the methods of Jahn *and others* (2016) and Zafar *and others* (2017), called SCITE and SiFit, respectively. SiFit uses an MCMC approach as a heuristic to find the maximum likelihood tree from imperfect SCS data. Based on the inferred tumor phylogenetic tree, SiFit estimates the mutation order by estimating the most likely mutation states of the tips and the internal nodes using a dynamic programming algorithm. Although both SCITE and SiFit by default output only the order of the mutations, both can be used to account for uncertainty in the inferred order. For example, because SCITE uses an MCMC algorithm for inference, the posterior probability associated with various mutation orders can be obtained by examining the frequency with which these orders are sampled by the MCMC algorithm. Similarly, the authors of SiFit recently developed a method called SiCloneFit (Zafar *and others*, 2019) that utilizes MCMC to sample trees, and thus the algorithm from SiFit for inferring mutation order on a fixed tree could be applied to a posterior sample of trees to measure the uncertainty in the mutation order that results from uncertainty in the tumor phylogeny.

In this paper, we propose a novel method for inferring the order in which mutations arise within an individual tumor given SCS data from the tumor at a single time point. Our approach utilizes models for both the mutational process within the tumor and the errors that arise during SCS data collection in a Bayesian framework, thus allowing us to quantify the uncertainty in the inferred mutation order along a fixed tumor phylogeny. Our approach thus represents a conceptually distinct and practically important extension of earlier methods.

## 2. Methods

We assume that we are given a phylogenetic tree with branch lengths that displays the evolutionary relationships among a sample of *J* cells within a tumor. To infer the locations (branches) on which a set of somatic mutations are acquired in the tree, we need to model the evolutionary process of the somatic mutations and quantify the technical errors that arise from the SCS data collection process. We assume that during the evolutionary process, somatic mutations evolve independently across sites, and each mutation evolves independently on different branches. We also assume that each somatic mutation occurs once along the phylogeny and that no back mutation occurs, so that all descendant cells linked by the mutation branch will harbor the corresponding mutation. When quantifying the effect of errors, we assume that SCS technical errors for mutations are independent of one another.

### 2.1 Notation and terminology

Consider somatic mutations of interest at *I* loci across the genome for a sample of *J* single cells. The *J* single cells are sampled from different spatial locations (clones) within the tumor. The mutation data can be either binary or ternary. For binary data, 0 denotes the absence of mutation and 1 means that mutation is present, while for ternary data, 0, 1 and 2 represent the homozygous reference (normal), heterozygous (mutation present) and homozygous non-reference (mutation present) genotypes, respectively.

The *I* somatic mutations evolve along the tumor evolutionary tree 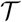. Each tip in 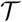 represents one single cell *C*_*j*_, where *j* = 1, … , *J*. Let *C* = {*C*_1_, … , *C*_*J*_} be the set of the *J* single cells under comparison. 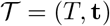 includes two parts: the tree topology *T* and a vector of branch lengths t. The tree topology *T* = (*V, E*) is a connected graph without cycles and is composed of nodes and branches, where *V* is the set of nodes and *E* is the set of branches. Trees are rooted, and the root *r* represents the common ancestor (a normal cell without somatic mutations) for all the single cells under comparison. In the context of this paper, all the definitions in the following sections will apply to rooted bifurcating trees. There are 2*J* − 2 branches in a rooted bifurcating tree with *J* tips, i.e., *E* = {*e*_1_, *e*_2_, … , *e*_2*J*−2_}. Let *v* and *w* be two nodes in the node set *V* that are connected by the branch *x* in the branch set *E* (i.e., *x* = {*v, w*}: *v* is the immediate ancestor node of *w*, and *x* connects *v* and *w*). Then the set *U*^*x*^(*w*), which includes the node *w* and all nodes descended from *w* in 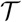, is called the *clade induced by w*. The branch *x* connects the ancestor node *v* and the clade induced by *w*, and we define branch *x* as the *ancestor branch of clade U*^*x*^(*w*). *E*^*x*^(*w*) is a subset of *E* that includes branches connecting nodes in *U*^*x*^(*w*), and *C*^*x*^(*w*) are the tips in *U*^*x*^(*w*).

Let *G*_*ij*_ denote the true genotype for the *i*^*th*^ genomic site of cell *C*_*j*_. The *i*^*th*^ genomic site will then have a vector G_*i*_ ∈ {0, 1}^*J*^ (for binary data) or {0, 1, 2}^*J*^ (for ternary data) representing its true genotype for all the *J* cells represented by the tips in the tree, where *i* = 1, … , *I*. Let *S*_*ij*_ denote the observed data for the *i*^*th*^ genomic site of cell *C*_*j*_. Due to the technical errors associated with SCS data, the observed data *S*_*ij*_ does not always equal the true genotype *G*_*ij*_. For both binary and ternary data, the observed state *S*_*ij*_ might be flipped with respect to the true genotype *G*_*ij*_ due to FP or FN. Missing states (“-”) or low-quality states (“?”) may be present for some genomic sites as well. Fig. 2 shows an example of true and observed binary genotype data for the mutations in Fig. 1. In Fig. 2, the observed state is highlighted in red if it is not consistent with the true genotype. The red numbers are those mutations with flipped observed mutation states relative to the true mutation states. The red dash (“-”) indicates a missing value and the red question mark (“?”) represents a low-quality value.

**Fig. 2:**
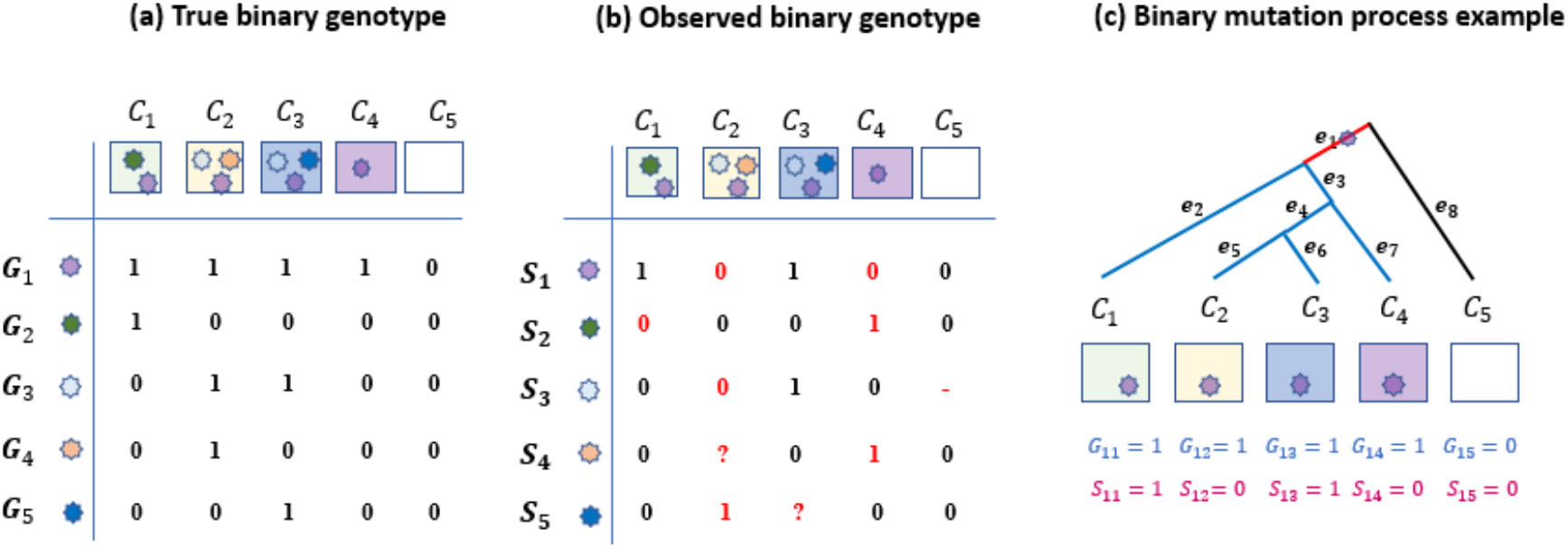
True binary data, observed binary data and binary mutation process example. (a) True binary mutation matrix of the sequenced tumor cells in the mutation tree in Fig. 1(e). Each row represents true genotypes for one genomic site in all cells and each column represents the true genotypes of multiple genomic sites for one single cell. (b) Observed mutation matrix with missing and ambiguous values (red), as well as mutation states that are misrecorded with respect to the true mutation matrix (red numbers; these are either false positives or false negatives). The red dash indicates a missing value since the sequencing process does not return signal at this site of this cell, and the red question mark represents an ambiguous value. Each row represents observed states for one genomic site in all cells and each column represents the observed states of multiple genomic sites for one single cell. (c) Binary mutation process example. A mutation is acquired on branch *e*_1_ (highlighted in red). The cell descending from branch *e*_8_ (highlighted in black) does not carry the mutation, while the cells descending from the blue branches carry the mutation.

Mathematically, we represent the observed mutation states of the *J* single cells at *I* different genomic sites by an *I* × *J* mutation matrix **S** for convenience,

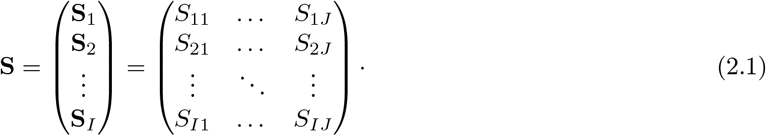

Each entry (*i, j*) denotes the state observed for mutation *i* in cell *C*_*j*_, so **S**_*i*_ gives the observed data for genomic site *i* as a vector with *J* values corresponding to the *J* single cells. Column *j* represents the mutations of interest for cell *C*_*j*_. In 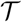, let 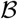 be the vector of locations (branches) on which the *I* mutations occur, i.e., 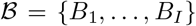, where *B*_*i*_ is the branch on which mutation *i* is acquired. Note that *B*_*i*_ takes values in {e_1_, e_2_, … , e_2*J*−2_}.

### 2.2 Somatic mutation process

To model the somatic mutation process, we consider continuous-time Markov processes, which we specify by assigning a rate to each possible transition between states. We consider point mutations. Once a mutation *i* is acquired on a branch *x* ∈ *E*, all the branches in the set *E*^*x*^(*w*) will harbor mutation *i* but those branches in the set *E*\(*x* ∪ *E*^*x*^(*w*)) will not carry this mutation.

#### 2.2.1 Binary genotype data

For binary genotype data, the mutation process can be modeled by the 2 × 2 transition rate matrix

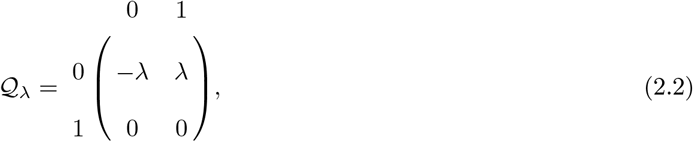

where *λ* denotes the instantaneous transition rate per genomic site. The transition probability matrix *P* (*t*) along a branch of length *t* is then computed by matrix exponentiation of the product of 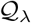 and the branch length *t*, which gives

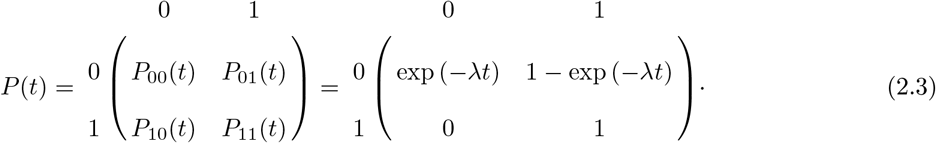

Note that *P*_01_(*t*) is the probability that mutation *i* is acquired along a branch of length *t*. Under this model and recalling that each mutation evolves independently along different branches in 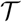, the marginal probability that mutation *i* is acquired on branch *x* ∈ *E*, denoted by 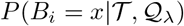 is thus given by

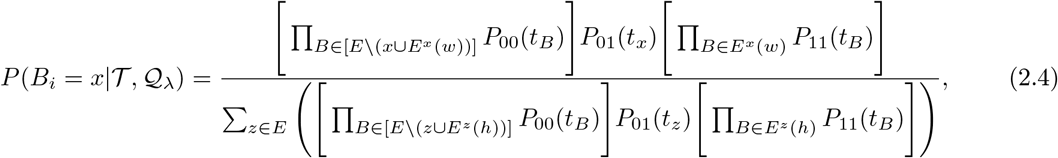

where *t*_*B*_ is length of branch *B*. In the numerator, the first term is a product of probabilities over all branches without the mutation, the second term is the probability that the mutation is acquired on branch *x*, and the third term is a product of probabilities over all branches with the mutation, i.e., all branches in *E*^*x*^(*w*). The denominator is needed to create a valid probability distribution over all possible branches, and is obtained by summing the numerator over all valid branches *z* ∈ *E*. The 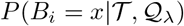 term is normalized by the denominator because we exclude two possibilities: a mutation is not acquired on any branch in 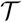, or a mutation is acquired more than once on different branches in 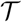.

As an example, Fig. 2 (c) depicts the observed and true binary genotype for mutation *i* = 1 shown in Fig. 2 (a)-(b). The set of branches is *E* = {*e*_1_, … , *e*_8_} and the corresponding set of branch lengths would be t = {*t*_1_, … , *t*_8_}. If mutation *i* is acquired on branch *e*_1_, the cell descending along branch *e*_8_ will not carry the mutation, while those descending from the blue branches would carry this mutation. The marginal probability that mutation *i* = 1 is acquired on branch *e*_1_ would be proportional to its numerator, i.e., 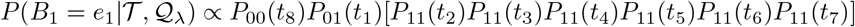.

#### 2.2.2 Ternary genotype data

The mutation model for ternary data is complex and includes three possible ways that mutation *i* occurs on a branch *x* in
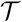:

1. The status of mutation *i* transitions from 0 → 1 on a branch *x* and there is no further mutation at this genomic site in 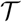.
2. The status of mutation *i* transitions directly from 0 → 2 on a branch *x* in 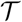.
3. The status of mutation *i* transitions from 0 → 1 on a branch *x* and then transitions from 1 → 2 on a branch *y* ∈ *E*^*x*^(*w*) in 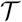.

We let *B*_*i*_ be the location at which mutation *i* occurs, 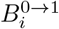 would be the branch on which mutation status transitions from 0 to 1, 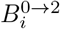 is the branch on which mutation status transitions from 0 to 2, and 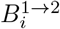 is the branch on which mutation status transitions from 1 to 2. If the mutation *i* occurs on branch *x*, all cells in *C*^*x*^(*w*) will carry 1 or 2 mutations. In other words, *G*_*ij*_ = 1 or 2 for all *C*_*j*_ ∈ *C*^*x*^(*w*) and *G*_*ij*_ = 0 for all *C*_*j*_ ∈ *C*\*C*^*x*^(*w*). We define the transition rate matrix 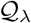 as

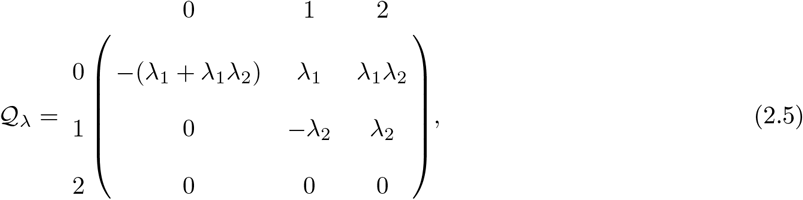

where *λ*_1_ and *λ*_2_ denote the instantaneous transition rates per genomic site of the transitions 0 → 1 and 1 → 2, respectively. Studies have provided evidence that direct mutation of 0 → 2 at rate *λ*_1_*λ*_2_ is possible in principle, although it is extremely rare (Iwasa *and others*, 2004). If *λ*_2_ is 0 in Expression (2.5), the model will be reduced to the infinite sites diploid model. The transition probability matrix 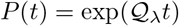 is then given by

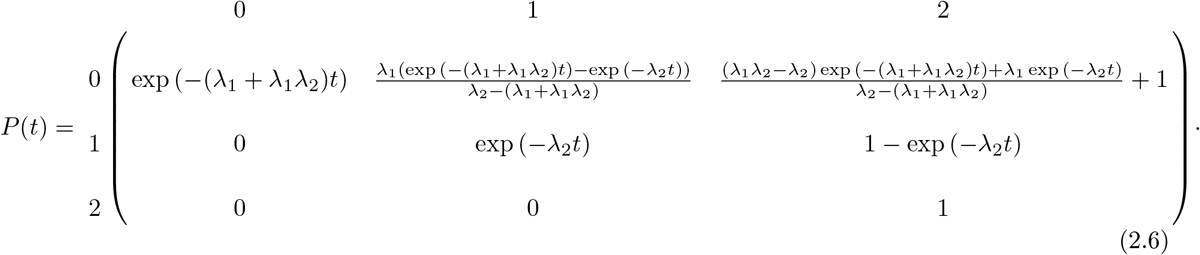

The marginal probability that mutation *i* occurs on branch *x* ∈ *E* for the three possible conditions is thus given by

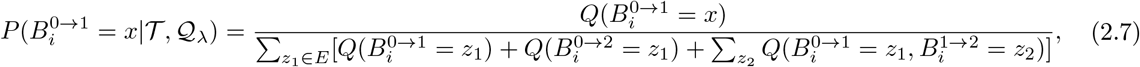

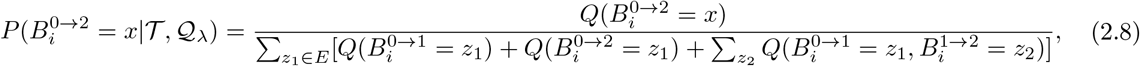

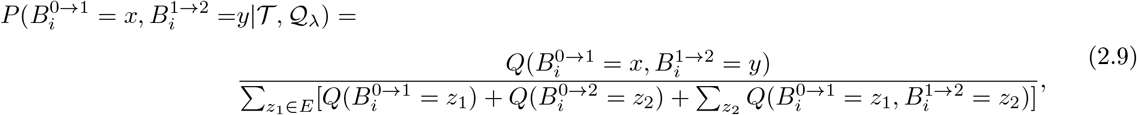

where

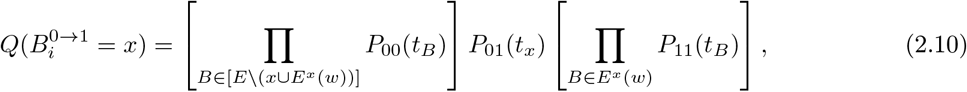

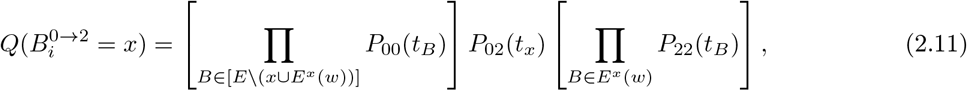

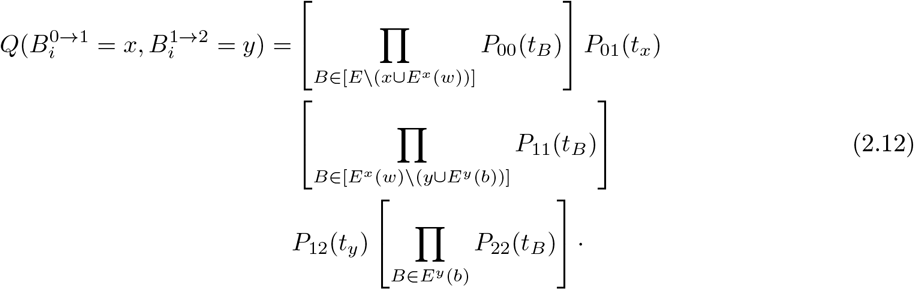

We normalize the marginal probabilities to exclude scenarios in which mutations are acquired more than once or in which mutations are not acquired in 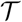. As an example, Fig. S1 in the Supplementary Material depicts the same mutation as in Fig. 2, but considers ternary data, leading to the following:

1. The marginal probability that mutation *i* transitions from 0 → 1 on branch *e*_1_ is 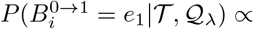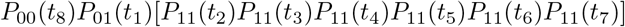.
2. The marginal probability that mutation *i* transitions from 0 → 2 on branch *e*_1_ is 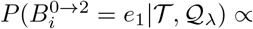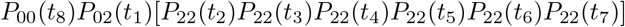.
3. The marginal probability that mutation *i* transitions from 0 → 1 on *e*_1_, and from 1 → 2 on *e*_3_ is 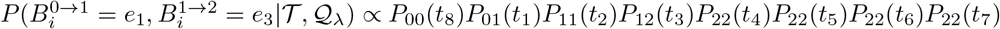.

The probability 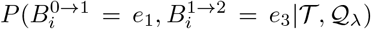 is the marginal probability that two mutations at the same site along the genome occur on two branches *e*_1_ and *e*_3_, respectively. After the first mutation occurs on branch *e*_1_, the second mutation can occur on any branch except *e*_1_ and *e*_8_.

### 2.3 Quantification of SCS errors

To account for FPs and FNs in the observed SCS data, our method applies the error model for binary and ternary data from Kim and Simon (2014), Jahn *and others* (2016), and Zafar *and others* (2017). Let *α*_*ij*_ be the probability of a false positive error and *β*_*ij*_ be the probability of a false negative error for genomic site *i* of cell *C*_*j*_.

For binary data, if the true genotype is 0, we may observe a 1, which is a false positive error. If the true genotype is 1, we may observe a 0, which is a false negative error. The conditional probabilities of the observed data given the true genotype at genomic site *i* of cell *C*_*j*_ are

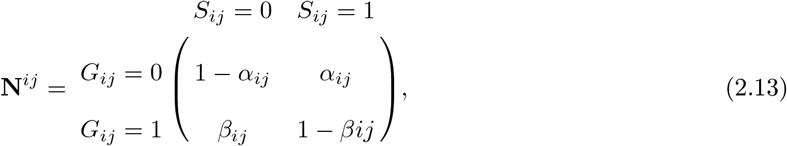

where 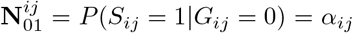, and other entries are defined similarly. Under the assumption that sequencing errors are independent, if mutation *i* is acquired on branch *x*, we can precisely quantify the effect of SCS technical errors for mutation *i* as

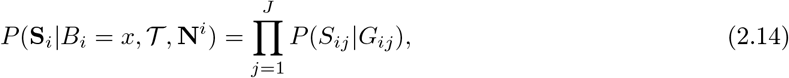

where N^*i*^ = {N^*i1*^, … , N^*iJ*^}. Using the example in Fig. 2, the error probability of the observed genotype conditioning on the mutation *i* = 1 occurring on branch *e*_1_ would be 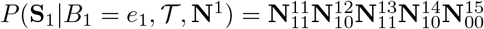, where N^1^ = {N^11^, … ,N^15^} for this binary data example.

For ternary data, the conditional probabilities of the observed data given the true genotype are given by

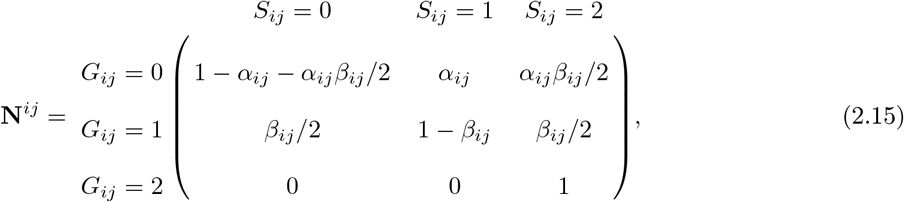

where 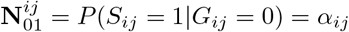, and the other entries are defined similarly. Under the same assumptions as for binary genotype data, we can precisely quantify the effect of SCS technical errors as in Equation (2.14) if mutation *i* is acquired on branch *x*. Using the example in Fig. S1 in the Supplementary Material, the error probabilities for the three possible ways that mutation *i* = 1 may arise on branch *e*_1_ are

1. The error probability under the condition that the true mutation transitions from 0 → 1 on branch *e*_1_ is 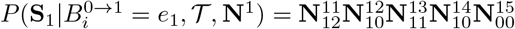.
2. The error probability under the condition that the true mutation transitions from 0 → 2 on branch *e*_1_ is 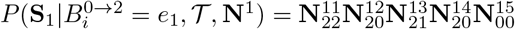.
3. The error probability under the condition that the true mutation transitions from 0 → 1 on branch *e*_1_, and transitions from 1 → 2 on branch *e*_3_ is 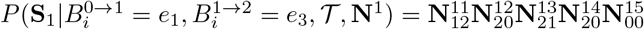

And N^1^ = {N^11^, … , N^15^} for this ternary data example. The term 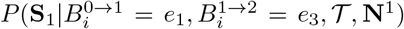 gives the error probability for the case in which the two mutations at the same genomic site occur on branches *e*_1_ and *e*_3_.

### 2.4 Missing and low-quality data

In real data, missing and low-quality states are observed and must be taken into account. For each mutation *i*, we exclude cells with missing states, and a subtree 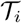 from 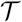 is extracted. The number of tips *J*_*i*_ in subtree 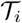 is less than or equal to *J*. Let *E*_*i*_ be the set of branches in subtree 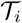. The probability that mutation *i* occurs on branch *x* is then given by 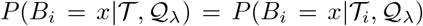, where 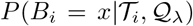 is computed based on branches in the subtree 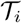, and 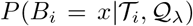 is 0 for those branches *x* ∈ *E*\*E*_*i*_. We quantify the effect of the SCS technical errors as

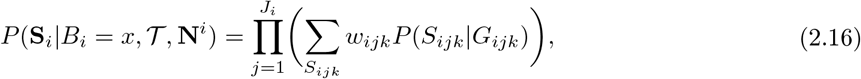

where *w*_*ijk*_ is the weight for each possible observed state at a mutation site. For a site with an observed state that is not missing or ambiguous, *w*_*ijk*_ is 1 for the observed state and 0 for all other states. For an ambiguous site, we can assign equal weight for each possible state, or we can assign weight based on sequencing information or other biological characteristics.

### 2.5 Inferring the location of a mutation in 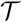

Once the observed matrix **S** = [**S**_1_ … **S**_*I*_]^*T*^ of the *I* mutations has been collected, the next step is to infer the branch on which mutation *i* takes place, conditioning on **S**. Given the observed data matrix **S**, the tumor phylogenetic tree 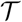, the error probability matrix N = {N^*ij*^ |1 ⩽ *i* ⩽ *I,* 1 ⩽ *j* ⩽ *J*}, and the mutation process 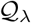, we can assign a posterior probability distribution 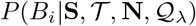 to the location of mutation *i* using Bayes’ theorem,

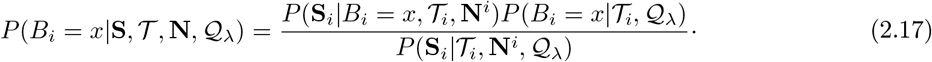

For mutation *i*, 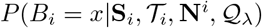 is computed for all *x* in set *E*_*i*_. For example, there are 8 branches in the tree in Fig. 2, so the branch on which mutation *i* = 1 occurs, *B*_1_, can be any of the 8 branches. For the binary example, the posterior probability that mutation *i* = 1 occurs on *e*_1_ is 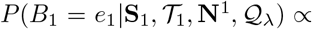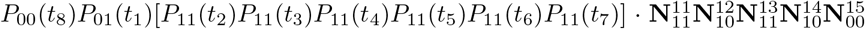. In this way, the posterior probability that the mutation occurs on each of the 8 branches can be computed, giving the probability distribution for the location of mutation *i* = 1, i.e. 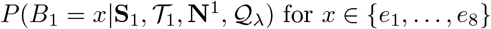.

To summarize this probability distribution, we construct a (1 − *θ*) × 100% credible set for the location of mutation *i* as follows. First, the branches are ranked by their posterior probabilities, and then branches are added to the credible set in the order of decreasing posterior probability until the sum of their probabilities reaches (1 − *θ*). The number of branches in the credible set is informative about the level of certainty associated with the inferred location for the mutation. To obtain a point estimate, we pick the branch that maximizes the posterior probability, i.e., the maximum a posteriori (MAP) estimate. The MAP estimator for the location of mutation *i* is given by

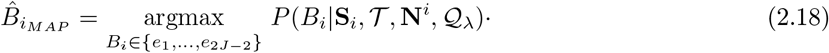

For the example in Fig. 2, the branch with the largest posterior probability is 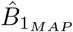 for mutation *i* = 1.

### 2.6 Inferring the mutation order in 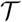

We now consider the joint posterior probability distribution of the locations for the *I* mutations in the sample of *J* single cells. Based on the assumption of independence among the *I* mutations being considered, the posterior distribution for 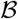 is given by

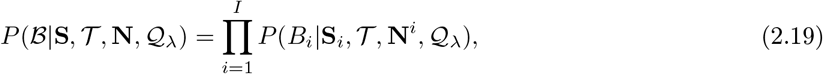

where N^*i*^ = {N^*i*1^, … , N^*iJ*^}. From this distribution, we can extract information on the ordering of mutations of interest. For example, if we are interested in the order of mutation *i* = 1 and mutation *i* = 2 in Fig. 2, the joint posterior probability distribution that mutation *i* = 1 occurs on branch *x* ∈ *E* and mutation *i* = 2 occurs on branch *y* ∈ *E* can be used to find the probability that mutation *i* = 1 occurs earlier in the tree than mutation *i* = 2. Note that 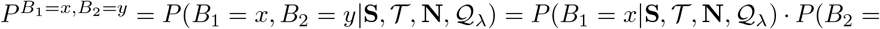 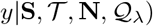. This joint distribution can be represented in a matrix given by

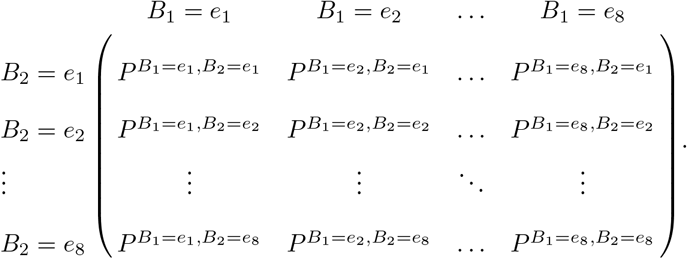

Adding entries of the matrix for which branch *B*_1_ is earlier in the tree than branch *B*_2_ thus gives the probability that mutation 1 occurs before mutation 2. To measure the uncertainty of the ordering of the mutations, we rank all possible mutation orders by their posterior probabilities, and construct a (1−*θ*)×100% credible set by adding orders with decreasing probability until the sum exceeds 1 − *θ*. The MAP estimator for the order of *I* mutations is thus given by

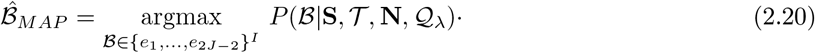

## 3. Simulation Study

To evaluate the ability of our method, which we call MO (Mutation Order), to correctly identify the locations and the order of a set of mutations under different conditions, we conduct a series of simulation studies with data simulated under different assumptions. The goal is to assess the effect of data quality (complete or incomplete, high or low error probabilities), number of cells, branch lengths, number of mutations and type of genotype data on the performance of our method. We consider a total of 12 scenarios, with 100 replicates for each setting within each scenario. Scenarios 1 - 4 involve data generated under our model for either 10 cells (scenarios 1 and 2) or 50 cells (scenarios 3 and 4) for either long branch lengths (scenarios 1 and 3) or short branch lengths (scenarios 2 and 4). Scenarios 5 - 8 consider data simulated under various models implemented in the CellCoal software (Posada, 2020). Scenarios 9 and 10 involve data generated under our model, but with mutations placed on branches with varying (rather than equal) probabilities. Finally, scenarios 11 and 12 consider data simulated under the finite sites assumption (all other simulation settings use the infinite sites assumption). The methods used to simulate data under these different scenarios are described in detail in Section A of the Supplementary Material, and Section D of the Supplementary Material provides information about computational requirements.

### 3.1 Accuracy of MAP estimates

We assess the accuracy of the MAP estimates in MO across the 100 trees within each simulation setting in several ways, including whether the mutation is inferred to occur on the correct branch (“location accuracy”), whether any pair of mutations are inferred to occur in the correct order (“order accuracy”), and whether a pair of mutations that occur on adjacent branches are inferred to occur in the correct order (“adjacent order accuracy”). In evaluating both the order accuracy and adjacent order accuracy, if two sequential mutations are inferred to occur on the same branch, then it is counted as ordering the mutations incorrectly. In addition, pairs of mutations that occur on the same branch are also included in the computation of order accuracy and adjacent order accuracy. The details of how the MAP estimates are assessed are given in Section B of the Supplementary Material. Tables 1 to 4 in the Supplementary Material show the location accuracy for scenarios 1 to 4 with each cell entry corresponding to a unique setting of *α*, *β*, type of genotype and missing data percentage. In most cases, the location accuracy of MO is high except when the error probabilities are high. The accuracy rates of settings with 50 cells in Tables 3 and 4 are slightly higher than those with 10 cells in Tables 1 and 2. With the same type of genotype and same error probability setting, the accuracy decreases as the percentage of missing values increases. When *α* (or *β*) is fixed, accuracy decreases as *β* (or *α*) increases.

The results for order accuracy (Tables 5 to 8 in the Supplementary Material) and adjacent order accuracy (Tables 9 to 12 in the Supplementary Material) for scenarios 1 to 4 are similar. In addition to the same overall trend due to number of cells, data type, percentage of missing data and error probabilities, the order accuracy rates are higher than the corresponding adjacent order accuracy rates.

The results for location accuracy, order accuracy and adjacent order accuracy of MO in scenarios 5 to 10 have similar patterns to those observed for scenarios 1 to 4. The accuracy in scenarios 5 to 10 is not affected by the number of mutations. In addition to the same overall trend due to the number of cells, type of genotype, missing data percentage and error probabilities, the accuracy rates in scenarios 5 to 10 are higher than the corresponding accuracy rates in scenarios 1 to 4. Especially when error probabilities are low, the accuracy can be as high as 99%.

### 3.2 Credible set accuracy

The credible set accuracy of the inferred mutation branch is assessed as well. If the true mutation branch is within the credible set, we count this as correct; otherwise, it is incorrect. We use 95% credible set for computation (Tables 13 to 16 for scenarios 1 to 4 in the Supplementary Material). The credible set accuracy has the same overall trend as the accuracy of MAP estimates due to the number of cells, type of genotype, missing data percentage and error probabilities, though the accuracy is much higher than that of the corresponding MAP estimates, especially for settings with large error probabilities and high missing data percentages. The overall trend for scenarios 5 to 10 is similar to scenarios 1 to 4, but the corresponding accuracy rates in scenarios 5 to 10 are higher than those in scenarios 1 to 4.

### 3.3 Comparison with competing approaches

To further assess the performance of MO, we compare its performance with the methods SCITE (Jahn *and others*, 2016) and SiFit (Zafar *and others*, 2017) for the simulation data in scenarios 1 to 12. SCITE can estimate the order of mutations for either binary or ternary genotype data. We use the maximum likelihood mutation order inferred by SCITE with 1,000,000 iterations given the true error probabilities. SiFit can use either binary or ternary genotype data when inferring the phylogenetic tree, but it can only use binary genotype data when inferring mutation order. We estimate the most likely mutational profiles for the tips and the internal nodes by SiFit given the true phylogenetic tree, error probabilities and mutation rates. We then extract the mutation order information from the output. The three methods are compared with respect to the order accuracy and adjacent order accuracy for the above simulation settings.

#### 3.3.1 Scenarios 1 to 4

Fig. 3 and Fig. 4 plot the order accuracy and the adjacent order accuracy for the three methods in scenarios 1 to 2, respectively. Fig. S2 and Fig. S3 in the Supplementary Material plot the order accuracy and the adjacent order accuracy for the three methods in scenarios 3 to 4, respectively. In each figure, the top row shows the results for binary data and the bottom row shows the results for ternary data. In each panel, different methods are highlighted in different colors.

**Fig. 3:**
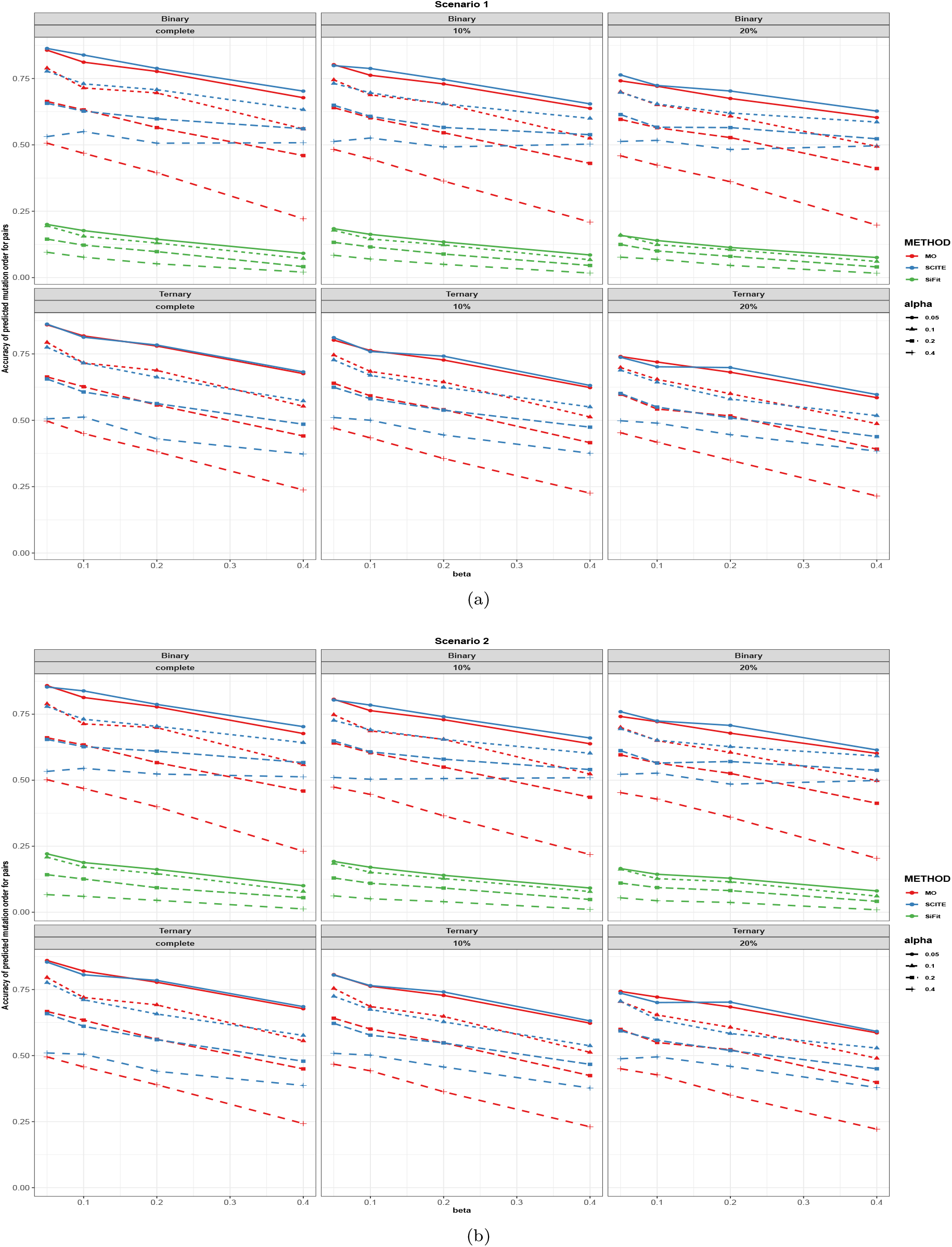
Order accuracy in scenarios 1 and 2 for MO, SCITE and SiFit. Each panel includes the results from the specific type of genotype and missing data percentage. In each panel, red, blue and green colors correspond to MO, SCITE and SiFit, respectively. Each plotting symbol on the line represents a different probability of a false positive error, *α*. The x-axis is the probability of a false negative error, *β*, and the y-axis is order accuracy.

**Fig. 4:**
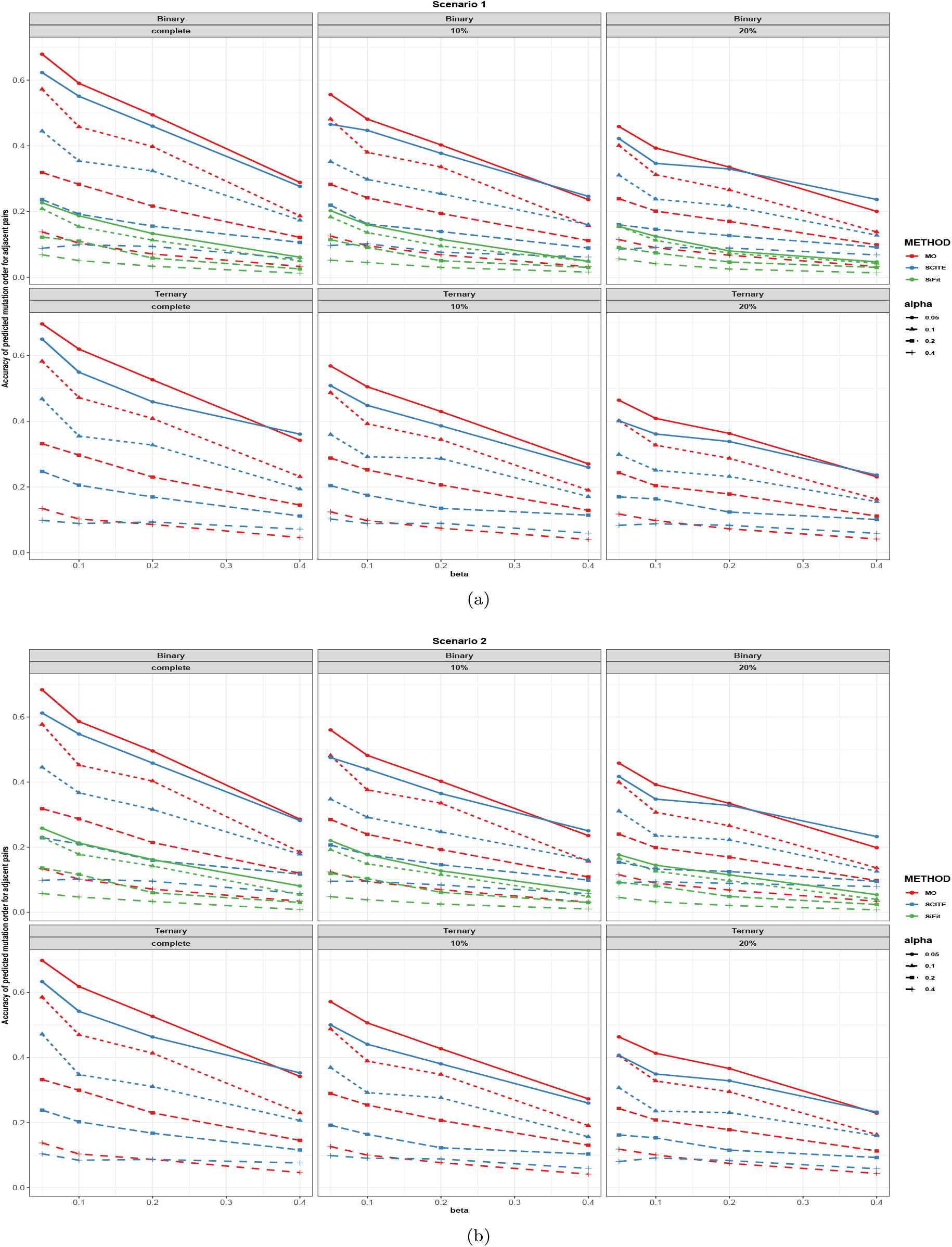
Adjacent order accuracy in scenarios 1 and 2 for MO, SCITE and SiFit. Each panel includes the results from the specific type of genotype and missing data percentage. In each panel, red, blue and green colors correspond to MO, SCITE and SiFit, respectively. Each plotting symbol on the line represents a different probability of a false positive error, *α*. The x-axis is the probability of a false negative error, *β*, and the y-axis is adjacent order accuracy.

In scenarios 1 to 4, order accuracy and adjacent order accuracy show decreasing trend as data quality becomes worse for all three methods. For results estimated from the trees with 10 cells (scenarios 1 and 2), MO has slightly higher adjacent order accuracy estimated from binary and ternary data. Only when both *α* and *β* are large does SCITE have higher adjacent order accuracy rates than MO. Comparing order accuracy when there are 10 cells in each tree, MO has comparable order accuracy when error probabilities are small. MO has lower order accuracy than SCITE when error probabilities are large but the discrepancies of order accuracy rates between MO and SCITE are only 5% on average. When there are 50 cells in each tree (scenarios 3 and 4), MO is superior to SCITE in all settings in terms of order accuracy and adjacent order accuracy estimated from both binary and ternary genotype data. Specifically, the order accuracy for MO is 25% higher than SCITE on average, and the adjacent order accuracy for MO is 20% higher than SCITE on average. In all settings, SiFit has the worst performance since only a subset of the input mutations are inferred to occur on the tree. Although the output partial mutation order from SiFit are mostly correct, the accuracy is low due to the small number of inferred mutation orders. MO thus dominates SiFit when assessing the performance using order accuracy and adjacent order accuracy. Comparing between settings with 10 cells and those with 50 cells, the performance of MO is consistently good, and the accuracy increases as the number of cells increases. SiFit performs better as the number of cells increases as well. However, the performance of SCITE becomes worse when the number of cells increases. Although the number of correct pairs inferred by SCITE increases, the accuracy decreases because the total number of true pairs increases.

#### 3.3.2 Scenarios 5 to 8

Fig. S4 and Fig. S5 in the Supplementary Material plot the order accuracy and adjacent order accuracy for scenarios 5 and 6, respectively. In scenarios 5 and 6 where mutations evolve by the infinite sites diploid model, order accuracy and adjacent order accuracy show decreasing trend as data quality becomes worse for all three methods, as is observed for scenarios 1 to 4. MO is superior to SCITE in all settings in terms of adjacent order accuracy and order accuracy. In all the settings, SiFit has the worst performance with respect to order accuracy. However, SiFit has comparable adjacent order accuracy to SCITE when error probabilities are small. Similar to scenarios 1 to 4, only a proportion of mutations are inferred to occur on the tree by SiFit. MO thus dominates SiFit in scenarios 5 and 6. In all settings for scenarios 5 and 6, the number of mutations and number of tips in the tree do not affect the order accuracy or adjacent order accuracy of MO and SiFit very much. However, the performance of SCITE is affected by the number of mutations. As the number of mutations increases, the accuracy of SCITE becomes lower. In addition, the adjacent order accuracy of SCITE increases as the number of cells increases.

Fig. S6 and Fig. S7 in the Supplementary Material plot the order accuracy and the adjacent order accuracy for scenarios 7 and 8, respectively. In scenarios 7 and 8, mutations arise by the infinite sites diploid model, as was the case for scenarios 5 and 6, but now a small proportion of the mutations are lost. Compared to the complete settings in scenarios 5 and 6, the performance of all the three methods becomes worse. However, the performance of the three methods is comparable to settings with missing values in scenarios 5 and 6.

In addition to the above comparisons, we also apply MO to data from scenarios 5 and 6 when transition rates are misspecified. Fig. S11 and Fig. S12 show the order accuracy and adjacent order accuracy when MO is applied with misspecified transition rates. In each panel, MO, SCITE, and SiFit are highlighted in red, blue, and green, respectively, when the transition rates *λ*_1_ = 1 and *λ*_2_ = 0 are used, as in the initial analysis in scenarios 5 and 6. Purple color corresponds to MO when the misspecified transition rates are used. The performance of SCITE is not affected by misspecified transition rates. Comparing the plots, we see that when binary data are used, the effect of misspecified transition rates are ignorable. However, when using ternary data, the differences are noticeable. In scenario 5, the order accuracy for MO with misspecified transition rates is comparable to SCITE when error probabilities are small and higher than SCITE when error probabilities are large. In scenario 6, the order accuracy inferred from ternary genotype data for MO with misspecified transition rates is lower than SCITE. Comparing the adjacent order accuracy with ternary data, the performance of MO with the misspecified transition rates is worse than when the transition rates are correctly specified in MO, but MO still performs better than SCITE.

#### 3.3.3 Scenarios 9 to 10

In scenarios 9 and 10, mutations are simulated under the mutation process defined in Section 2.2. Although the transition rates are the same as in scenarios 1 to 4, each mutation is not equally likely to occur on all of the branches. In Fig. S8 and Fig. S9, we observe that MO has higher accuracy than SCITE and SiFit in all settings in terms of both order accuracy and adjacent order accuracy.

#### 3.3.4 Scenarios 11 to 12

In scenarios 11 and 12, mutations are simulated under the finite sites assumption. Because it is unclear how mutation order should be defined when mutations can arise multiple times along a phylogeny, we instead plot the location accuracy of MO and SiFit in Fig. S10. When there are only 10 tips in the tree, most simulated mutations occur only once along the tree and MO has higher accuracy than SiFit. However, when there are 50 tips, most are back mutations and/or parallel mutations. SiFit performs better than MO when the rate of mutating from 1 to 0 is low. When the rate of mutating from 1 to 0 is high, neither MO nor SiFit identify the correct mutation location. MO is limited by its assumption that all mutations occur only once on the tree. Although SiFit can infer parallel/back mutations, it is not able to identify all the locations on which the mutations occur for the simulated data.

## 4. Empirical examples

We apply MO to two experimental single-cell DNA sequencing datasets, one for prostate cancer (Su *and others*, 2018) and one for metastatic colorectal cancer patients (Leung *and others*, 2017). For the prostate cancer dataset, we retrieve publicly available data from the single-cell study of Su *and others* (2018), which includes 10 single-cell genomes for each patient. For the colorectal cancer dataset, we use the somatic single nucleotide variants (SNVs) after variant calling provided in the original study (16 SNVs for patient CRC1 and 36 SNVs for patient CRC2) of Leung *and others* (2017).

### 4.1 Prostate cancer data

#### 4.1.1 Data analysis

To infer tumor evolutionary trees for patients 1 and 2 (labeled P1 and P2), we use the SVDQuartets method of Chifman and Kubatko (2014) as implemented in PAUP* (Swofford, 1999) using the aligned DNA sequences for all somatic mutations as input with the expected rank of the flattening matrix set to 4. We specify the normal cell sample as the outgroup. We use the maximum likelihood method to estimate the branch lengths.

We select common tumor suppressor genes and oncogenes for both P1 and P2 identified by Su *and others* (2018). In addition to these common cancer-associated genes across different cancers, we map mutations in prostate cancer-specific genes (genes that are more commonly mutated in prostate cancer patients) suggested by Barbieri *and others* (2013) and Tate *and others* (2018). For both binary and ternary genotype data for these genes, we use MO to compute the posterior probability of mutation on each branch of the tumor phylogeny for each of the two patients. Su *and others* (2018) estimated the error probabilities to be (*α, β*) = (0·29, 0·02) for P1, and (*α, β*) = (0·31, 0·02) for P2. Although our method in Section 2 allows the assignment of varying error probabilities across genomic sites and cells, here we use same probabilities for all sites.

To examine the effect of informativeness of the prior distribution on the resulting inference, we consider two prior distributions for each parameter with mean equal to the estimated error probability from the empirical data and with either a large or a small variance as described in Section C in the Supplementary Material. For P1, we consider *α*|**S**_*i*_ ~ *Beta*(0·29, 0·71) (larger variance) and *α*|**S**_*i*_ ~ *Beta*(2·9, 7·1) (smaller variance). For P2, we consider *α*|**S**_*i*_ ~ *Beta*(0·31, 0·69) (larger variance) and *α*|**S**_*i*_ ~ *Beta*(3·1, 6·9) (smaller variance). For *β* for both P1 and P2, we consider *β*|**S**_*i*_ ~ *Beta*(0·02, 0·98) (larger variance) and *β*|**S**_*i*_ ~ *Beta*(0·2, 9·8) (smaller variance).

According to Iwasa *and others* (2004), the mutation rates for the first and second mutation are estimated to be *λ*_1_ = 10^−7^ and *λ*_2_ = 10^−2^, respectively. We use these values to specify the prior distributions for the transition rates. Similar to the sequencing error probabilities, we set two prior distributions for each transition rate with equal means but different variances. The distribution of the transition rate *λ*_1_ (0 → 1 for ternary genotype) is set as *λ*_1_|**S**_*i*_ ~ *Gamma*(2, 5·0 × 10^−8^) (larger variance) and *λ*_1_|**S**_*i*_ ~ *Gamma*(5, 2·0 × 10^−8^) (smaller variance). The distribution of the transition rate *λ*_2_ (1 → 2 for ternary genotype) is set as *λ*_2_|**S**_*i*_ ~ *Gamma*(2, 5·0 × 10^−3^) (larger variance) and *λ*_2_|**S**_*i*_ ~ *Gamma*(5, 2·0 × 10^−3^) (smaller variance). The estimated probabilities of mutation do not vary substantially when the prior distributions with larger or smaller variance are used for any of these parameters. The heatmaps of estimated probabilities with different prior distributions (larger or smaller variance) are in the Supplementary Material.

#### 4.1.2 Results

Fig. 5 and Fig. S13 show the tumor evolutionary tree estimated for P1 and P2, respectively. In both tumor trees, the trunk connects the tumor clone to the normal clone. We annotate the genes on their inferred mutation branches. The uncertainty in the inferred mutation locations is highlighted in colors. Mutations with strong signal (defined to be a posterior probability larger than 0.7 that the mutation occurred on a single branch) are colored red, while mutations with moderate signal (defined to be a total posterior probability larger than 0.7 on two or three branches) are colored blue. Note that the posterior probability on a branch measures the support in the data under the model and prior distribution for the placement of the mutation on that branch. Mutations colored red are those for which the placement on a single branch is strongly supported. Mutations colored blue are those for which there is strong support for the mutation having occurred on one of the indicated branches. This should not be interpreted as evidence that the mutation occurred more than once; rather, it means that the precise placement of the mutation is ambiguous but can be confidently limited to the branches indicated.

**Fig. 5:**
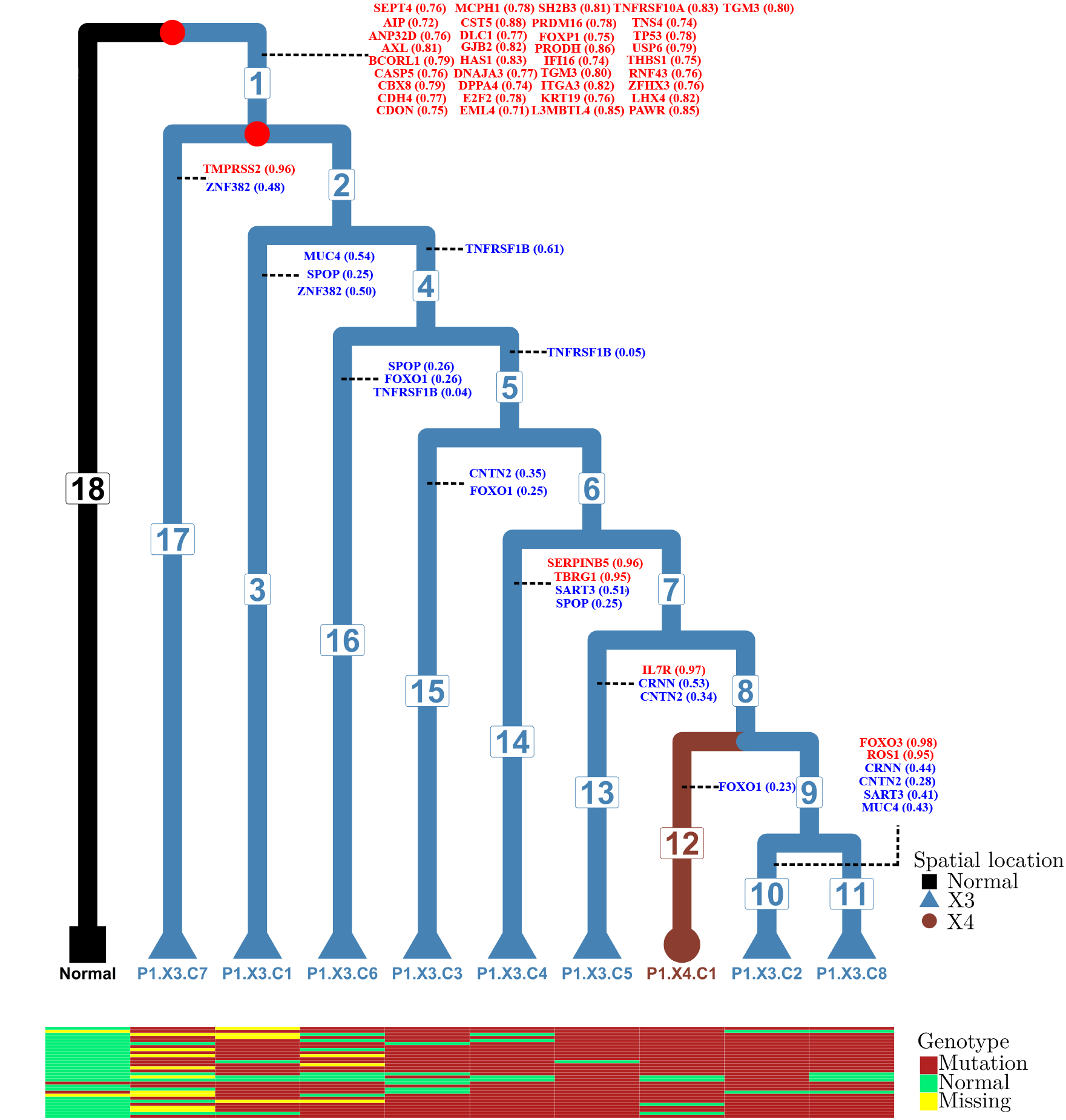
P1 tumor phylogenetic tree and inferred temporal order of the mutations. The normal cell is set as the outgroup. There are 18 branches in this tree. We do not assume the molecular clock when estimating the branch lengths. Branch lengths in this figure are not drawn to scale. The color and tip shape represent the spatial locations of the samples (normal tissue, location X3 or location X4; see Su *and others* (2018)). The temporal order of the mutations is annotated on the branches of the tree. Mutations with very strong signals (probability of occurring on one branch is greater than 0.7) are highlighted in red, while mutations with moderate signals (probabilities that sum to more than 0.7 on two or three branches) are highlighted in blue. Mutation data for 30 genes corresponding to the first 30 rows in Fig. S16 and Fig. S17 for each tip are shown in the heatmap matrix at the bottom.

We also compare the estimated posterior probability distributions for mutations of common cancer-associated genes for patients P1 and P2, which are used to construct credible sets and to measure the uncertainty of the inferred mutation order. Fig. S16 to Fig. S19 are the posterior probability distributionheatmaps for patients P1 and P2 with different prior distributions (larger or smaller variance).

Fig. S20 to Fig. S23 show heatmaps of the estimated posterior probabilities for prostate cancer-specific genes for patients P1 and P2 with different prior distributions (larger or smaller variance). In agreement with the results of Su et al. (2018), we find that mutation of *TP53*, which is commonly associated with tumor initiation in many cancers (see, e.g., Yu *and others* (2014)), is inferred to occur on the trunk of the tree with high probability in patient P1, but not on the trunk of the tumor tree of patient P2. Gene *ZFHX3* has a high probability of having mutated on the trunk of the tree in both patients. In addition, the heatmap for patient P1 shows strong signal that *FOXP1* mutates on the trunk of the tumor tree, while *BRCA2* has a high probability of having mutated on the trunk of the tree for patient P2. Comparing the heatmaps of common cancer-associated genes with the prostate cancer-specific genes, mutations inferred to have occurred on the trunk of the tree tend to be those that are common across cancer types, while mutations known to have high frequency within prostate cancer are generally found closer to the tips of the tree in both patients.

### 4.2 Metastatic colorectal cancer data

#### 4.2.1 Data analysis

The original study of Leung *and others* (2017) reported 16 and 36 SNVs for patients CRC1 and CRC2 after variant calling. We use SiFit (Zafar *and others*, 2017) to estimate each colorectal patient’s tumor phylogeny, including branch lengths, since there are fewer data available to estimate the phylogenies in this case and this method is specifically designed to estimate tumor phylogenies from single-cell data. The normal cells in each patient are merged into one normal sample and used as the outgroup.

Leung *and others* (2017) reported error probabilities of (*α, β*) = (0·0152, 0·0789) and (*α, β*) = (0·0174, 0·1256) for CRC1 and CRC2, respectively. For each patient, we use these values to specify the same prior distributions across all sites. For CRC1, we consider *α*|**S**_*i*_ ~ *Beta*(0·0152, 0·9848) (larger variance) and *α*|**S**_*i*_ ~ *Beta*(0·15, 9·85) (smaller variance); and *β*|**s**_*i*_ ~ *Beta*(0·078, 0·922) (larger variance) and *β*|**S**_*i*_ ~ *Beta*(0·78, 9·22) (smaller variance). For CRC2, we consider *α*|**S**_*i*_ ~ *Beta*(0·0174, 0·9826) (larger variance) and *α*|**S**_*i*_ ~ *Beta*(0·174, 9·826) (smaller variance); and *β*|**S**_*i*_ ~ *Beta*(0·1256, 0·8744) (larger variance) and *β*|**S**_*i*_ ~ *Beta*(1·256, 8·744) (smaller variance). The prior distributions for the transition rates for CRC1 and CRC2 are estimated by SiFit. As was found for the prostate cancer patients, the estimated probabilities do not vary substantially when we use prior distributions with small or large variance.

#### 4.2.2 Results

The inferred tumor trees and mutation order are depicted in Fig. S14 and Fig. S15 in the Supplementary Material. The posterior probabilities of the inferred mutation locations are indicated with colors as for the prostate cancer data, and agree overall with the findings of Leung *and others* (2017). Fig. S24 and Fig. S25 are heatmaps for the posterior probability distribution of each mutation for patients CRC1 and CRC2 with different priors. For patient CRC1, mutations in *APC*, *KRAS* and *TP53* are inferred to have been acquired on the trunk of the tumor phylogeny with high posterior probability, in agreement with Leung *and others* (2017) and in agreement with past studies. The studies of Fearon and Vogelstein (1990) and Powell *and others* (1992) have shown that the mutation order of these genes appears to be fixed in initializing colorectal cancer, providing further support for our findings. In addition, we identify the five mutations specific to metastatic cells that are found by Leung *and others* (2017), with three (*ZNF521*, *TRRAP*, *EYS*) inferred to occur on branch 97 in Fig. S14. Support is found for placement of *RBFOX1* and *GATA1* in two distinct regions of the tree. Each supported placement is on a branch that leads to a clade of metastatic aneuploid cells, indicating the association of such cells with these mutations. If the tree is correct, then this might indicate that these mutations arose more than once (though our model assumes that each mutation only arose once, if the “truth” is that the mutation arose more than once, a reasonable behavior of our model would be to partition the posterior probability between the two origins). Another possibility is that the tree is incorrect, and the two clades of metastatic aneuploid cells should actually be clustered together, which would then presumably result in strong support for the placement of these mutations on the branch leading to the combined clade.

For CRC2, we identify strong signals on branch 36 in Fig. S15 for several genes reported by Leung *and others* (2017) that are shared by primary and metastatic cells, including driver mutations in *APC*, *NRAS*, *CDK4* and *TP53*. We also identify an independent lineage of primary diploid cells (colored in pink in Fig. S15) that evolved in parallel with the rest of the tumor with moderate to strong signals for mutations in *ALK*, *ATR*, *EPHB6*, *NR3C2* and *SPEN* and that do not share the mutations listed in the previous sentence. Our analysis further agrees with that of Leung *and others* (2017) in that we also identify the subsequent formation of independent metastatic lineages. For example, on branches 56 and 58 we find moderate support for mutations in *FUS*; and strong support for mutation on branch 136 in *HELZ* and branch 78 in *PRKCB*. Many of the genes showing weaker or moderate support for mutation in these metastatic lineages agree with those identified by Leung *and others* (2017). The primary difference between our result and that of Leung *and others* (2017) is that we identify mutation in *NR4A3* and *FUS* to have occurred along a different metastatic lineage than the mutations in *TSHZ3* and *PRKCB*.

## 5. Discussion

Development of computational tools based on a phylogenetic framework for use in studying cancer evolution has the potential to provide tremendous insight into the mechanisms that lead to ITH, especially the role of the temporal order of mutations in cancer progression. For example, Ortmann et al. (2015) have shown differences in clinical features and the response to treatment for patients with different mutation orders, indicating that inference of the order in which mutations arise within an individual’s tumor may have direct implications in clinical oncology, for both diagnostic applications in measuring the extent of ITH and targeted therapy. SCS data provide an unprecedented opportunity to estimate mutation order at the highest resolution. However, such data are subject to extensive technical errors that arise during the process of whole-genome amplification.

To analyze such data, we introduce MO, a new Bayesian approach for reconstructing the ordering of mutational events from the imperfect mutation profiles of single cells. MO is designed to infer the temporal order of a collection of mutations of interest based on a phylogeny of cell lineages that allows modeling of the errors at each tip. MO can infer the mutation order that best fits single-cell data sets that are subject to the technical noise, including ADO, false positive errors, low-quality data, and missing data. The assumption of independence of mutations made by MO is the same as that made in other methods developed for inferring mutation order (e.g., Zafar *and others* (2017), Zafar *and others* (2019), and Jahn *and others* (2016)). Thus, MO does not presently account for possible interactions between the occurrences of mutations, though it could be extended to accommodate this if biological information about these interactions is available. However, recent work (Canisius *and others*, 2016) indicates that observed dependence typically takes the form of mutual exclusivity (i.e., only one gene in the group will be mutated in any given patient) rather than positive association, making the independence assumption of less concern here, as the set of mutations we study are assumed to be present within an individual patient. MO could also be extended to work on clonal trees and models that include errors in observed data for multiple cells in a tip instead of a single cell. In addition, MO could be modified to account for the accelerated mutation rates common in late-stage cancers, or to allow for back or parallel mutation.

An important difference between MO and existing methods, such as SCITE (Jahn *and others*, 2016) and SiFit (Zafar *and others*, 2017), is the mechanism for quantifying uncertainty in the inferred order. Options available within SCITE (Jahn *and others*, 2016) allow for estimation of the posterior probability distribution across orders. SiFit (Zafar *and others*, 2017), on the other hand, could be modified to account for uncertainty in the orders because the true tumor phylogeny is unknown and must first be estimated. In contrast, because MO uses a probabilistic model for inferring mutation locations along a fixed tree, it is able to provide an estimate of uncertainty in the inferred locations conditioning on the correct tumor phylogeny, thus capturing a source of uncertainty that differs from what SCITE and SiFit provide. MO performs accurately, as is evident from a comprehensive set of simulation studies that take into account different aspects of modern SCS data sets by examining a wide range of error probabilities, fractions of missing data, branch lengths, and numbers of cells in each tree. The simulation studies also demonstrate that MO outperforms the state-of-the-art methods when the number of cells is large and performs comparably to other methods when the number of cells is small. MO is robust to the technical errors that arise during whole-genome amplification. When applied to data from prostate cancer patients and colorectal cancer patients, MO is able to not only provide insight into the locations of cancer-associated mutations, but also the level of certainty in the locations. However, MO does not provide estimates of transition rates and error probabilities as SiFit and SCITE do, but rather integrates over uncertainty in these parameters.

The methodology underlying MO could be enhanced by incorporating models for copy number alterations, as well as by considering mutations that affect the same allele more than once. As SCS data collection becomes more advanced, enabling hundreds of cells to be analyzed in parallel at reduced cost and increased throughput, MO is poised to analyze the resulting large-scale data sets to make meaningful inference of the mutation order during tumor progression for individual patients. MO thus represents an important step forward in understanding the role of mutation order in cancer evolution and as such may have important translational applications for improving cancer diagnosis, treatment, and personalized therapy. If inferred mutation order can be associated with clinical outcomes, future research can explore the cause of clinical outcomes given specific mutation order with the goal of developing novel targeted treatments. This will allow clinical providers to make decisions concerning treatment based on the mutation landscapes of patients. Although the current study focuses on cancer, MO can potentially also be applied to single-cell mutation profiles from a wide variety of fields. These applications are expected to provide new insights into our understanding of cancer and other human diseases.

## 6. Software

MO has been implemented in R and is available at https://github.com/lkubatko/MO.

## 7. Supplementary Material

Supplementary material is available.

## 8. Acknowledgments

The simulation experiments and data analyses were carried out using the ASC Unity Cluster at The Ohio State University, USA. The authors thank two anonymous reviewers for helpful comments on an earlier draft of this manuscript.

