## Supplementary Material for "A Phylogenetic Approach to Inferring the Order in Which Mutations Arise during Cancer Progression"

Yuan Gao*[1], Jeff Gaither[2], Julia Chifman[3], and Laura Kubatko[4,5,6]

[1]Division of Biostatistics, The Ohio State University, Columbus, OH 43210

[2]Institute for Genomic Medicine, Nationwide Children's Hospital, Columbus, OH 43205

[3]Department of Mathematics and Statistics, American University, Washington, DC 20016

[4]Mathematical Biosciences Institute, The Ohio State University, Columbus, OH 43210

[5]Department of Statistics, The Ohio State University, Columbus, OH 43210

[6]Department of Evolution, Ecology, and Organismal Biology, The Ohio State University, Columbus, OH 43210

### Supplementary Figures

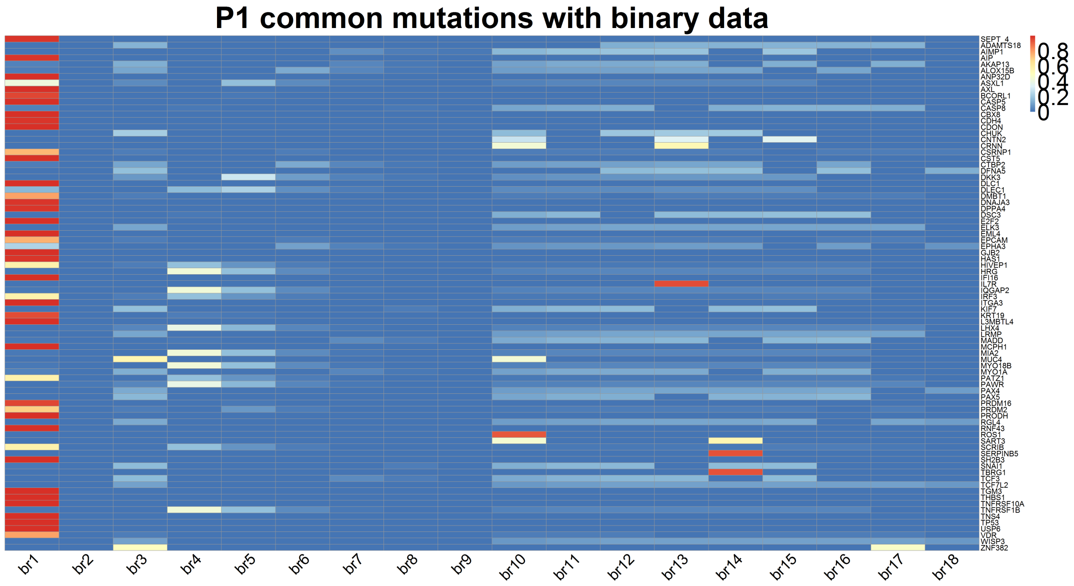


(a)

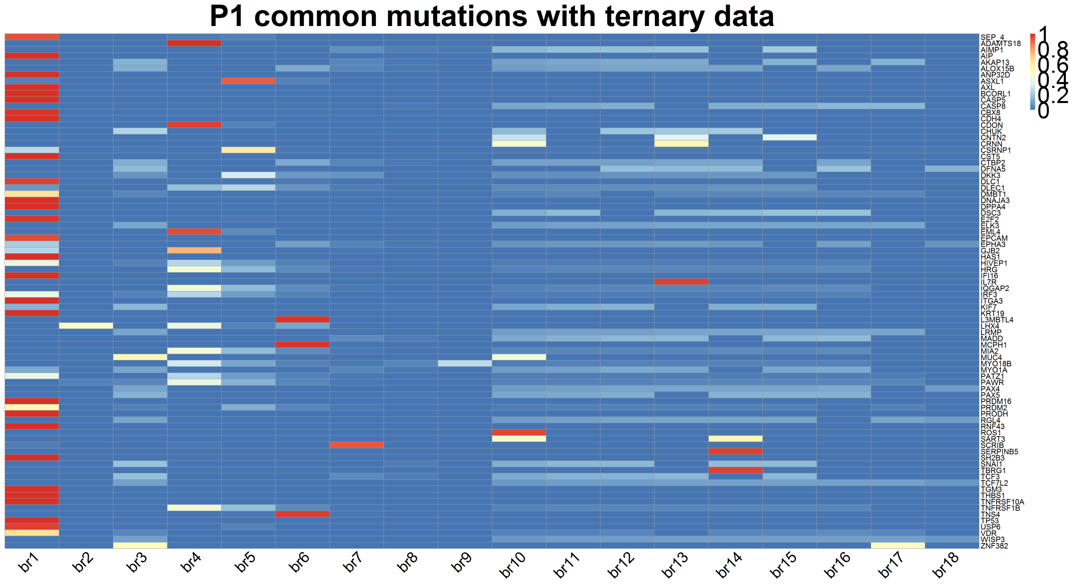


(b)

Figure S1: Heatmap of posterior probabilities that each mutation occurs on each branch for common tumor suppressor genes or oncogenes for prostate cancer patient P1 using either binary (a) or ternary (b) data. Colors indicate the magnitude of the probability, with red indicating probability close to 1 and blue indicating probability close to 0. For P1, prior of $\alpha$ is set as $\alpha|\mathbf{S}_i \sim Beta(0.29, 0.71)$ (larger variance). The prior of $\beta$ is set as $\beta|\mathbf{S}_i \sim Beta(0.02, 0.98)$ (larger variance). Distribution of mutation rate $\lambda_1$ ($0 \rightarrow 1$ for ternary genotype) is set as $\lambda_1|\mathbf{S}_i \sim Gamma(10^{-5}, 10^{-2})$ (larger variance). Distribution of mutation rate $\lambda_2$ ($1 \rightarrow 2$ for ternary genotype) is set as $\lambda_2|\mathbf{S}_i \sim Gamma(10^{-1}, 10^{-1})$ (larger variance).

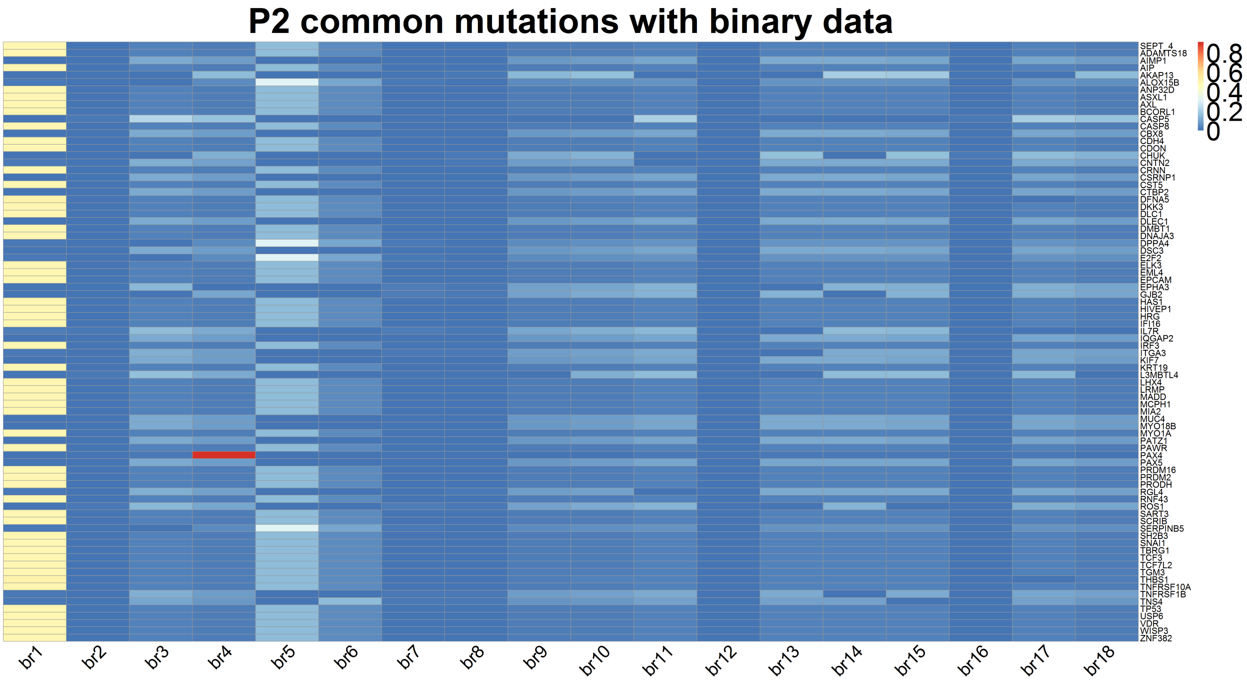


(a)

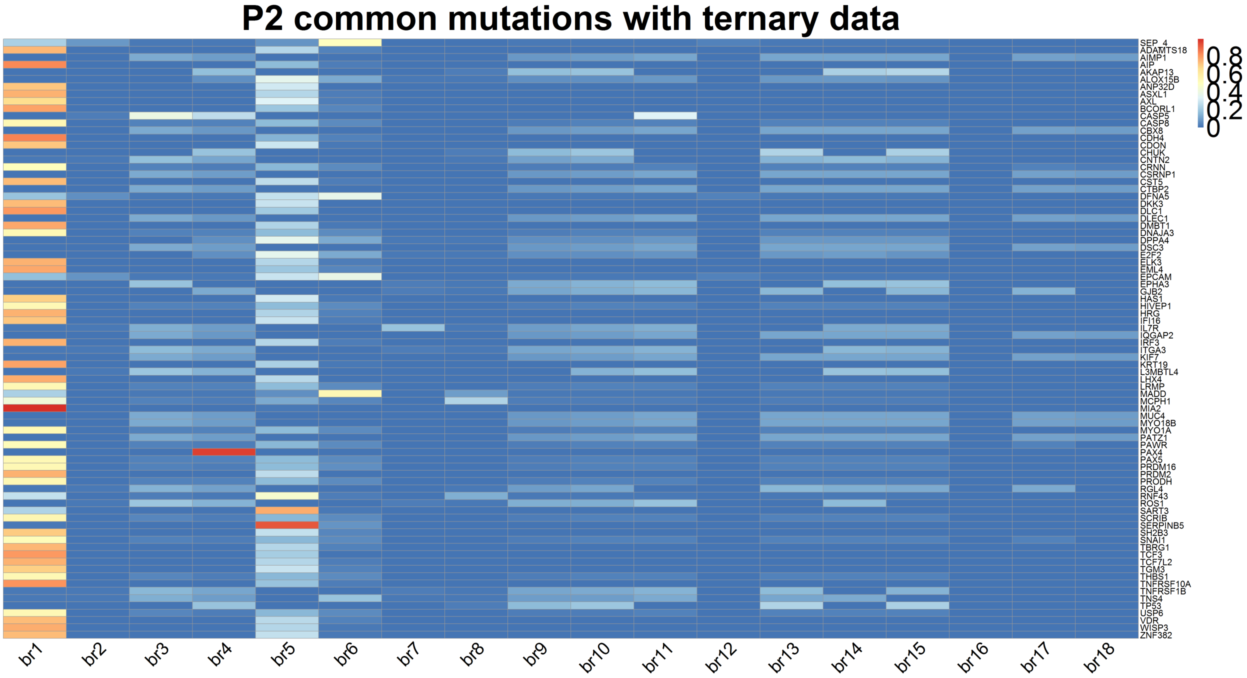


(b)

Figure S2: Heatmap of posterior probabilities that each mutation occurs on each branch for common tumor suppressor genes or oncogenes for prostate cancer patient P2 using either binary (a) or ternary (b) data. Colors indicate the magnitude of the probability, with red indicating probability close to 1 and blue indicating probability close to 0. For P2, prior distributions for the mutation rate parameters are as in Figure S1. Prior of $\alpha$ is set as $\alpha|\mathbf{S}_i \sim Beta(0.31, 0.69)$ (larger variance). The prior of $\beta$ is set as $\beta|\mathbf{S}_i \sim Beta(0.02, 0.98)$ (larger variance).

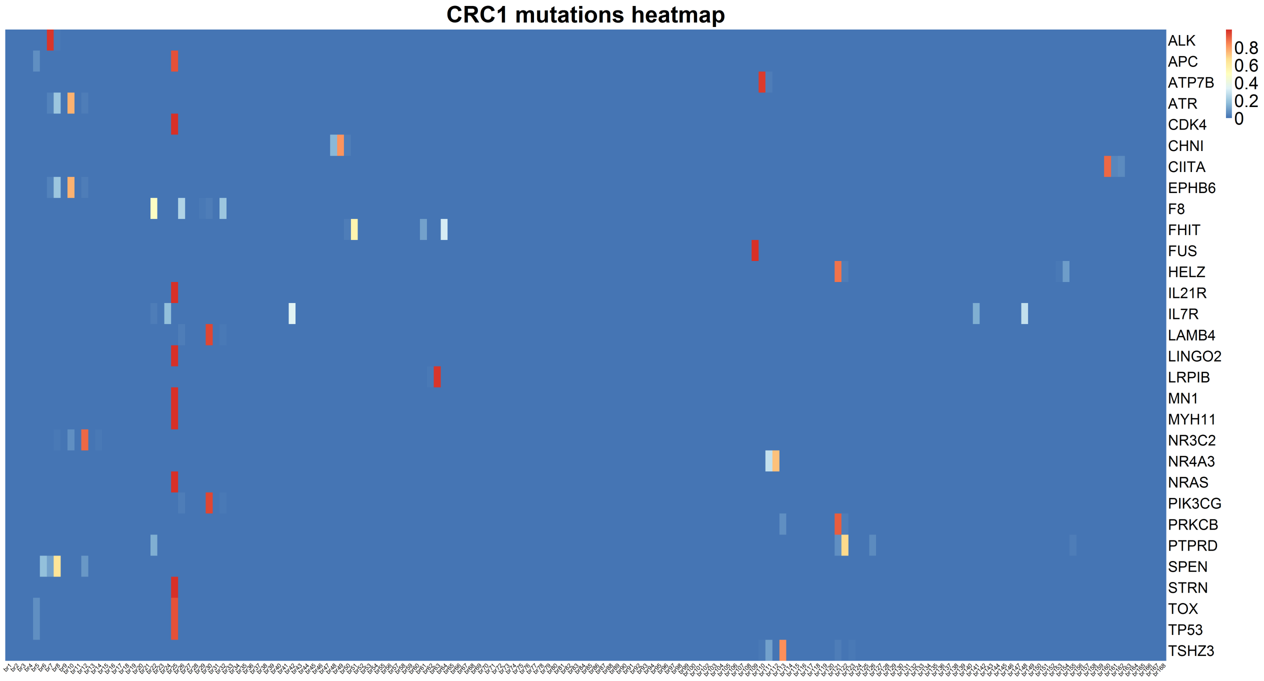


(a)

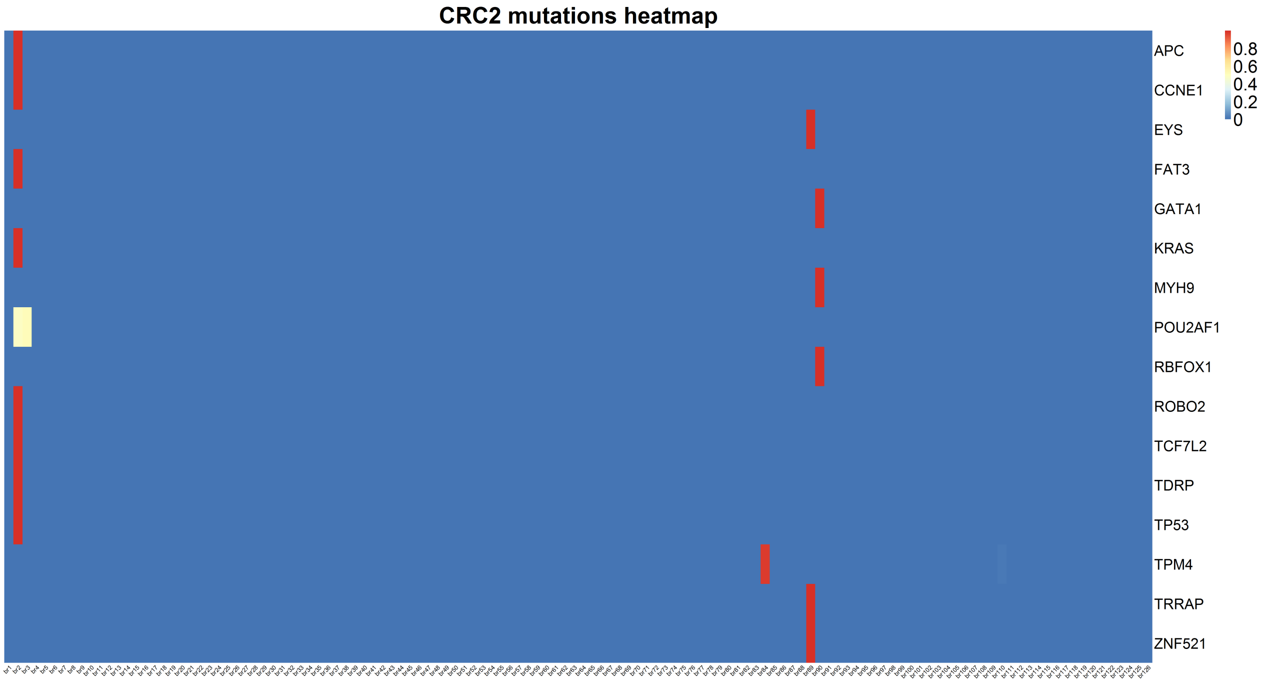


(b)

Figure S3: Heatmap of probabilities on each branch for mutations in patient CRC1 (a) and CRC2 (b) using binary data. Red color indicates very large mutation probability on that branch, while blue color indicates fairly small mutation probability. Mutation rates distributions are set as the same as P1.

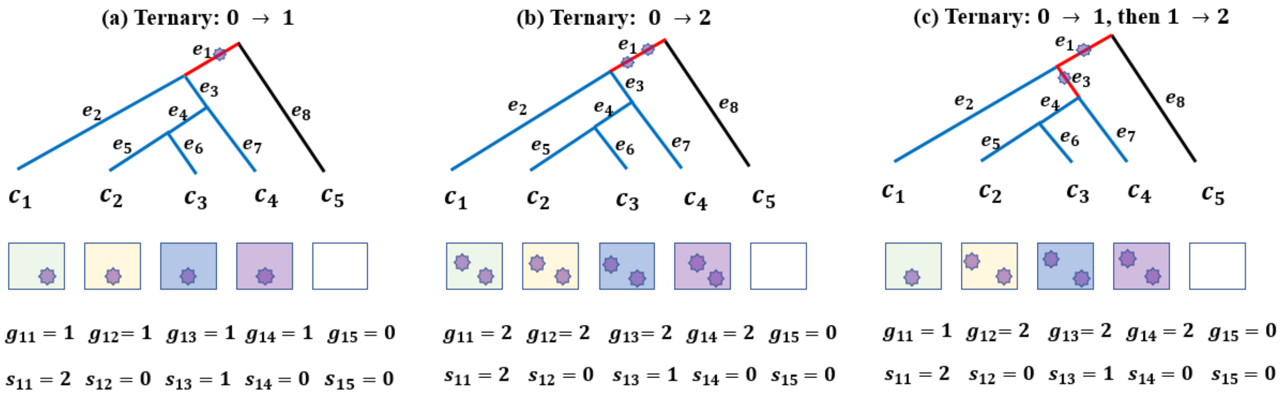


Figure S4: Three possible ways that a mutation may arise on branch $e_1$.

### Supplementary Tables

Table S1: Location accuracy of MO for scenario 1 where tree branch lengths follow an exponential distribution with mean 0.2. Each cell corresponds to unique $\alpha$ and $\beta$, type of genotype and missing data percentage (Columns 3-4: data with no missing values, columns 5-6: data with 10% of missing values, columns 7-8: data with 20% of missing values).

| **FPR** $\alpha$ | **FNR** $\beta$ | **Accuracy** ternary | **Accuracy** binary | **Accuracy** 10% ternary | **Accuracy** 10% binary | **Accuracy** 20% ternary | **Accuracy** 20% binary |
|---|---|---|---|---|---|---|---|
| 0.05 | 0.05 | 0.9225 | 0.9129 | 0.8181 | 0.8104 | 0.7200 | 0.7129 |
| 0.05 | 0.1 | 0.9108 | 0.8876 | 0.8097 | 0.7893 | 0.7086 | 0.6912 |
| 0.05 | 0.2 | 0.8084 | 0.7795 | 0.7168 | 0.6915 | 0.6297 | 0.6079 |
| 0.05 | 0.4 | 0.6893 | 0.6300 | 0.6134 | 0.5619 | 0.5388 | 0.4958 |
| 0.1 | 0.05 | 0.8872 | 0.8765 | 0.7901 | 0.7816 | 0.6926 | 0.6851 |
| 0.1 | 0.1 | 0.8438 | 0.8238 | 0.7541 | 0.7362 | 0.6617 | 0.6468 |
| 0.1 | 0.2 | 0.7772 | 0.7428 | 0.6938 | 0.6624 | 0.6113 | 0.5851 |
| 0.1 | 0.4 | 0.6342 | 0.5758 | 0.5661 | 0.5144 | 0.5014 | 0.4555 |
| 0.2 | 0.05 | 0.8146 | 0.8028 | 0.7307 | 0.7210 | 0.6449 | 0.6365 |
| 0.2 | 0.1 | 0.7748 | 0.7526 | 0.6936 | 0.6737 | 0.6107 | 0.5956 |
| 0.2 | 0.2 | 0.6894 | 0.6491 | 0.6175 | 0.5828 | 0.5481 | 0.5191 |
| 0.2 | 0.4 | 0.5381 | 0.4624 | 0.4872 | 0.4182 | 0.4349 | 0.3743 |
| 0.4 | 0.05 | 0.6822 | 0.6629 | 0.6156 | 0.5993 | 0.5492 | 0.5353 |
| 0.4 | 0.1 | 0.6225 | 0.5928 | 0.5639 | 0.5384 | 0.5045 | 0.4819 |
| 0.4 | 0.2 | 0.5228 | 0.4680 | 0.4739 | 0.4238 | 0.4292 | 0.3839 |
| 0.4 | 0.4 | 0.3444 | 0.2465 | 0.3177 | 0.2285 | 0.2918 | 0.2098 |

Table S2: Location accuracy of MO for scenario 2 where tree branch lengths follow an exponential distribution with mean 0.1. Each cell corresponds to unique $\alpha$ and $\beta$, type of genotype and missing data percentage (Columns 3-4: data with no missing values, columns 5-6: data with 10% of missing values, columns 7-8: data with 20% of missing values).

| **FPR** $\alpha$ | **FNR** $\beta$ | **Accuracy** ternary | **Accuracy** binary | **Accuracy** 10% ternary | **Accuracy** 10% binary | **Accuracy** 20% ternary | **Accuracy** 20% binary |
|---|---|---|---|---|---|---|---|
| 0.05 | 0.05 | 0.9239 | 0.9140 | 0.8238 | 0.8159 | 0.7226 | 0.7145 |
| 0.05 | 0.1 | 0.8828 | 0.8630 | 0.7852 | 0.7668 | 0.6864 | 0.6714 |
| 0.05 | 0.2 | 0.8106 | 0.7803 | 0.7210 | 0.6943 | 0.6331 | 0.6106 |
| 0.05 | 0.4 | 0.6889 | 0.6320 | 0.6128 | 0.5633 | 0.5387 | 0.4958 |
| 0.1 | 0.05 | 0.8889 | 0.8784 | 0.7907 | 0.7828 | 0.6923 | 0.6851 |
| 0.1 | 0.1 | 0.8453 | 0.8258 | 0.7539 | 0.7371 | 0.6624 | 0.6488 |
| 0.1 | 0.2 | 0.7672 | 0.7333 | 0.6835 | 0.6534 | 0.6016 | 0.5771 |
| 0.1 | 0.4 | 0.6336 | 0.5758 | 0.5653 | 0.5147 | 0.5012 | 0.4566 |
| 0.2 | 0.05 | 0.8179 | 0.8058 | 0.7319 | 0.7214 | 0.6446 | 0.6352 |
| 0.2 | 0.1 | 0.7726 | 0.7518 | 0.6927 | 0.6732 | 0.6096 | 0.5937 |
| 0.2 | 0.2 | 0.6879 | 0.6487 | 0.6168 | 0.5819 | 0.5486 | 0.5175 |
| 0.2 | 0.4 | 0.5441 | 0.4667 | 0.4873 | 0.4195 | 0.4356 | 0.3749 |
| 0.4 | 0.05 | 0.6839 | 0.6671 | 0.6188 | 0.6028 | 0.5528 | 0.5391 |
| 0.4 | 0.1 | 0.6233 | 0.5935 | 0.5648 | 0.5368 | 0.5063 | 0.4816 |
| 0.4 | 0.2 | 0.5266 | 0.4705 | 0.4789 | 0.4279 | 0.4315 | 0.3864 |
| 0.4 | 0.4 | 0.3413 | 0.2470 | 0.3151 | 0.2283 | 0.2860 | 0.2095 |

Table S3: Adjacent order accuracy of MO for scenario 1. Tree branch lengths follow an exponential distribution with mean 0.2. Each cell corresponds to unique $\alpha$ and $\beta$, type of genotype and missing data percentage (Columns 3-4: data with no missing values, columns 5-6: 10% of data missing, columns 7-8: 20% of data missing).

| **FPR** $\alpha$ | **FNR** $\beta$ | **Accuracy** ternary | **Accuracy** binary | **Accuracy** 10% ternary | **Accuracy** 10% binary | **Accuracy** 20% ternary | **Accuracy** 20% binary |
|---|---|---|---|---|---|---|---|
| 0.05 | 0.05 | 0.8294 | 0.8192 | 0.6738 | 0.6661 | 0.5348 | 0.5267 |
| 0.05 | 0.1 | 0.8148 | 0.7789 | 0.6597 | 0.6307 | 0.5173 | 0.4948 |
| 0.05 | 0.2 | 0.6264 | 0.5937 | 0.5067 | 0.4799 | 0.4000 | 0.3787 |
| 0.05 | 0.4 | 0.4523 | 0.3847 | 0.3681 | 0.3139 | 0.2953 | 0.2535 |
| 0.1 | 0.05 | 0.7567 | 0.7470 | 0.6146 | 0.6070 | 0.4833 | 0.4779 |
| 0.1 | 0.1 | 0.6811 | 0.6584 | 0.5547 | 0.5342 | 0.4353 | 0.4195 |
| 0.1 | 0.2 | 0.5678 | 0.5302 | 0.4644 | 0.4309 | 0.3721 | 0.3451 |
| 0.1 | 0.4 | 0.3668 | 0.3178 | 0.3036 | 0.2616 | 0.2466 | 0.2129 |
| 0.2 | 0.05 | 0.6106 | 0.6017 | 0.5025 | 0.4955 | 0.3993 | 0.3944 |
| 0.2 | 0.1 | 0.5458 | 0.5275 | 0.4478 | 0.4313 | 0.3570 | 0.3461 |
| 0.2 | 0.2 | 0.4259 | 0.3922 | 0.3496 | 0.3226 | 0.2830 | 0.2619 |
| 0.2 | 0.4 | 0.2540 | 0.2539 | 0.2128 | 0.1682 | 0.1770 | 0.1404 |
| 0.4 | 0.05 | 0.3832 | 0.3707 | 0.3217 | 0.3112 | 0.2631 | 0.2533 |
| 0.4 | 0.1 | 0.3202 | 0.3033 | 0.2694 | 0.2559 | 0.2235 | 0.2118 |
| 0.4 | 0.2 | 0.2230 | 0.1940 | 0.1877 | 0.1619 | 0.1595 | 0.1358 |
| 0.4 | 0.4 | 0.0963 | 0.0575 | 0.0862 | 0.0543 | 0.0779 | 0.0463 |

Table S4: Adjacent order accuracy of MO for scenario 2. Tree branch lengths follow an exponential distribution with mean 0.1. Each cell corresponds to unique $\alpha$ and $\beta$, type of genotype and missing data percentage (Columns 3-4: data with no missing values, columns 5-6: 10% of data missing, columns 7-8: 20% of data missing).

| **FPR** $\alpha$ | **FNR** $\beta$ | **Accuracy** ternary | **Accuracy** binary | **Accuracy** 10% ternary | **Accuracy** 10% binary | **Accuracy** 20% ternary | **Accuracy** 20% binary |
|---|---|---|---|---|---|---|---|
| 0.05 | 0.05 | 0.8334 | 0.8334 | 0.6811 | 0.6734 | 0.5406 | 0.5322 |
| 0.05 | 0.1 | 0.7554 | 0.7280 | 0.6137 | 0.5884 | 0.4798 | 0.4619 |
| 0.05 | 0.2 | 0.6293 | 0.5909 | 0.5077 | 0.4762 | 0.4030 | 0.3787 |
| 0.05 | 0.4 | 0.4457 | 0.3889 | 0.3611 | 0.3162 | 0.2916 | 0.2551 |
| 0.1 | 0.05 | 0.7569 | 0.7468 | 0.6142 | 0.6074 | 0.4800 | 0.4756 |
| 0.1 | 0.1 | 0.6758 | 0.6552 | 0.5508 | 0.5339 | 0.4363 | 0.4226 |
| 0.1 | 0.2 | 0.5562 | 0.5197 | 0.4501 | 0.4187 | 0.3559 | 0.3333 |
| 0.1 | 0.4 | 0.3694 | 0.3204 | 0.3017 | 0.2622 | 0.2458 | 0.2157 |
| 0.2 | 0.05 | 0.6169 | 0.6076 | 0.5050 | 0.4965 | 0.4001 | 0.3928 |
| 0.2 | 0.1 | 0.5427 | 0.5241 | 0.4457 | 0.4282 | 0.3539 | 0.3408 |
| 0.2 | 0.2 | 0.4224 | 0.3927 | 0.3489 | 0.3223 | 0.2871 | 0.2646 |
| 0.2 | 0.4 | 0.2623 | 0.2056 | 0.2152 | 0.1683 | 0.1765 | 0.1386 |
| 0.4 | 0.05 | 0.3868 | 0.3761 | 0.3228 | 0.3119 | 0.2669 | 0.2577 |
| 0.4 | 0.1 | 0.3201 | 0.3022 | 0.2713 | 0.2544 | 0.2243 | 0.2119 |
| 0.4 | 0.2 | 0.2232 | 0.1928 | 0.1883 | 0.1609 | 0.1635 | 0.1368 |
| 0.4 | 0.4 | 0.0967 | 0.0579 | 0.0874 | 0.0516 | 0.0781 | 0.0486 |

Table S5: Order accuracy of MO for scenario 1. Tree branch lengths follow an exponential distribution with mean 0.2. Each cell corresponds to unique $\alpha$ and $\beta$, type of genotype and missing data percentage (Columns 3-4: data with no missing values, columns 5-6: 10% of data missing, columns 7-8: 20% of data missing).

| **FPR** $\alpha$ | **FNR** $\beta$ | **Accuracy** ternary | **Accuracy** binary | **Accuracy** 10% ternary | **Accuracy** 10% binary | **Accuracy** 20% ternary | **Accuracy** 20% binary |
|---|---|---|---|---|---|---|---|
| 0.05 | 0.05 | 0.9292 | 0.9281 | 0.8541 | 0.8548 | 0.7834 | 0.7830 |
| 0.05 | 0.1 | 0.9257 | 0.9183 | 0.8525 | 0.8463 | 0.7726 | 0.7672 |
| 0.05 | 0.2 | 0.8311 | 0.8337 | 0.7614 | 0.7650 | 0.6906 | 0.6935 |
| 0.05 | 0.4 | 0.7351 | 0.7298 | 0.6718 | 0.6686 | 0.6064 | 0.6059 |
| 0.1 | 0.05 | 0.8934 | 0.8927 | 0.8211 | 0.8216 | 0.7442 | 0.7460 |
| 0.1 | 0.1 | 0.8499 | 0.8505 | 0.7845 | 0.7842 | 0.7100 | 0.7103 |
| 0.1 | 0.2 | 0.7888 | 0.7900 | 0.7266 | 0.7279 | 0.6598 | 0.6614 |
| 0.1 | 0.4 | 0.6580 | 0.6680 | 0.6053 | 0.6123 | 0.5453 | 0.5523 |
| 0.2 | 0.05 | 0.8145 | 0.8151 | 0.7528 | 0.7546 | 0.6854 | 0.6868 |
| 0.2 | 0.1 | 0.7779 | 0.7767 | 0.7174 | 0.7180 | 0.6478 | 0.6513 |
| 0.2 | 0.2 | 0.6906 | 0.6956 | 0.6333 | 0.6414 | 0.5788 | 0.5853 |
| 0.2 | 0.4 | 0.5337 | 0.5397 | 0.4941 | 0.4978 | 0.4474 | 0.4475 |
| 0.4 | 0.05 | 0.6617 | 0.6589 | 0.6099 | 0.6080 | 0.5565 | 0.5540 |
| 0.4 | 0.1 | 0.5986 | 0.5995 | 0.5493 | 0.5532 | 0.4992 | 0.5007 |
| 0.4 | 0.2 | 0.4892 | 0.4906 | 0.4503 | 0.4504 | 0.4133 | 0.4165 |
| 0.4 | 0.4 | 0.3042 | 0.2813 | 0.2888 | 0.2612 | 0.2702 | 0.2407 |

Table S6: Order accuracy of MO for scenario 2. Tree branch lengths follow an exponential distribution with mean 0.1. Each cell corresponds to unique $\alpha$ and $\beta$, type of genotype and missing data percentage (Columns 3-4: data with no missing values, columns 5-6: 10% of data missing, columns 7-8: 20% of data missing).

| **FPR** $\alpha$ | **FNR** $\beta$ | **Accuracy** ternary | **Accuracy** binary | **Accuracy** 10% ternary | **Accuracy** 10% binary | **Accuracy** 20% ternary | **Accuracy** 20% binary |
|---|---|---|---|---|---|---|---|
| 0.05 | 0.05 | 0.9296 | 0.9275 | 0.8615 | 0.8607 | 0.7803 | 0.7793 |
| 0.05 | 0.1 | 0.8964 | 0.8945 | 0.8261 | 0.8229 | 0.7461 | 0.7445 |
| 0.05 | 0.2 | 0.8330 | 0.8339 | 0.7644 | 0.7658 | 0.6947 | 0.6961 |
| 0.05 | 0.4 | 0.7279 | 0.7265 | 0.6675 | 0.6694 | 0.6044 | 0.6062 |
| 0.1 | 0.05 | 0.8949 | 0.8937 | 0.8227 | 0.8238 | 0.7453 | 0.7468 |
| 0.1 | 0.1 | 0.8546 | 0.8534 | 0.7866 | 0.7865 | 0.7128 | 0.7153 |
| 0.1 | 0.2 | 0.7831 | 0.7841 | 0.7211 | 0.7217 | 0.6514 | 0.6559 |
| 0.1 | 0.4 | 0.6596 | 0.6673 | 0.6039 | 0.6130 | 0.5495 | 0.5560 |
| 0.2 | 0.05 | 0.8186 | 0.8190 | 0.7552 | 0.7559 | 0.6827 | 0.6826 |
| 0.2 | 0.1 | 0.7743 | 0.7752 | 0.7119 | 0.7134 | 0.6450 | 0.6480 |
| 0.2 | 0.2 | 0.6862 | 0.6935 | 0.6319 | 0.6373 | 0.5788 | 0.5833 |
| 0.2 | 0.4 | 0.5510 | 0.5469 | 0.5036 | 0.5028 | 0.4574 | 0.4562 |
| 0.4 | 0.05 | 0.6655 | 0.6631 | 0.6152 | 0.6105 | 0.5627 | 0.5593 |
| 0.4 | 0.1 | 0.6008 | 0.5994 | 0.5519 | 0.5511 | 0.5034 | 0.5017 |
| 0.4 | 0.2 | 0.4900 | 0.4895 | 0.4527 | 0.4505 | 0.4152 | 0.4141 |
| 0.4 | 0.4 | 0.3070 | 0.2835 | 0.2862 | 0.2589 | 0.2662 | 0.2406 |

Table S7: Credible set accuracy of MO for scenario 1. Tree branch lengths follow an exponential distribution with mean 0.2. Each cell corresponds to unique $\alpha$ and $\beta$, type of genotype and missing data percentage (Columns 3-4: data with no missing values, columns 5-6: 10% of data missing, columns 7-8: 20% of data missing).

| **FPR** $\alpha$ | **FNR** $\beta$ | **Accuracy** ternary | **Accuracy** binary | **Accuracy** 10% ternary | **Accuracy** 10% binary | **Accuracy** 20% ternary | **Accuracy** 20% binary |
|---|---|---|---|---|---|---|---|
| 0.05 | 0.05 | 0.9820 | 0.9833 | 0.9166 | 0.9186 | 0.8503 | 0.8528 |
| 0.05 | 0.1 | 0.9845 | 0.9866 | 0.9224 | 0.9251 | 0.8550 | 0.8594 |
| 0.05 | 0.2 | 0.9656 | 0.9720 | 0.9041 | 0.9113 | 0.8406 | 0.8481 |
| 0.05 | 0.4 | 0.9501 | 0.9570 | 0.8923 | 0.8992 | 0.8296 | 0.8362 |
| 0.1 | 0.05 | 0.9772 | 0.9804 | 0.9154 | 0.9196 | 0.8507 | 0.8554 |
| 0.1 | 0.1 | 0.9726 | 0.9777 | 0.9115 | 0.9179 | 0.8469 | 0.8544 |
| 0.1 | 0.2 | 0.9629 | 0.9683 | 0.9036 | 0.9098 | 0.8416 | 0.8486 |
| 0.1 | 0.4 | 0.9463 | 0.9515 | 0.8891 | 0.8932 | 0.8288 | 0.8315 |
| 0.2 | 0.05 | 0.9737 | 0.9780 | 0.9123 | 0.9173 | 0.8474 | 0.8534 |
| 0.2 | 0.1 | 0.9683 | 0.9736 | 0.9094 | 0.9143 | 0.8466 | 0.8517 |
| 0.2 | 0.2 | 0.9593 | 0.9633 | 0.9005 | 0.9049 | 0.8371 | 0.8420 |
| 0.2 | 0.4 | 0.9346 | 0.9326 | 0.8784 | 0.8749 | 0.8161 | 0.8146 |
| 0.4 | 0.05 | 0.9634 | 0.9685 | 0.9046 | 0.9089 | 0.8393 | 0.8447 |
| 0.4 | 0.1 | 0.9597 | 0.9587 | 0.9018 | 0.9022 | 0.8378 | 0.8390 |
| 0.4 | 0.2 | 0.9407 | 0.9382 | 0.8841 | 0.8803 | 0.8228 | 0.8196 |
| 0.4 | 0.4 | 0.8991 | 0.8720 | 0.8436 | 0.8179 | 0.7878 | 0.7607 |

Table S8: Credible set accuracy of MO for scenario 2. Tree branch lengths follow an exponential distribution with mean 0.1. Each cell corresponds to unique $\alpha$ and $\beta$, type of genotype and missing data percentage (Columns 3-4: data with no missing values, columns 5-6: 10% of data missing, columns 7-8: 20% of data missing).

| **FPR** $\alpha$ | **FNR** $\beta$ | **Accuracy** ternary | **Accuracy** binary | **Accuracy** 10% ternary | **Accuracy** 10% binary | **Accuracy** 20% ternary | **Accuracy** 20% binary |
|---|---|---|---|---|---|---|---|
| 0.05 | 0.05 | 0.9821 | 0.9842 | 0.9186 | 0.9216 | 0.8502 | 0.8542 |
| 0.05 | 0.1 | 0.9732 | 0.9784 | 0.9119 | 0.9175 | 0.8454 | 0.8523 |
| 0.05 | 0.2 | 0.9627 | 0.9698 | 0.9023 | 0.9101 | 0.8392 | 0.8473 |
| 0.05 | 0.4 | 0.9497 | 0.9560 | 0.8900 | 0.8972 | 0.8274 | 0.8353 |
| 0.1 | 0.05 | 0.9761 | 0.9797 | 0.9149 | 0.9192 | 0.8480 | 0.8536 |
| 0.1 | 0.1 | 0.9709 | 0.9766 | 0.9103 | 0.9170 | 0.8455 | 0.8521 |
| 0.1 | 0.2 | 0.9613 | 0.9680 | 0.9014 | 0.9090 | 0.8374 | 0.8455 |
| 0.1 | 0.4 | 0.9436 | 0.9499 | 0.8862 | 0.8916 | 0.8262 | 0.8297 |
| 0.2 | 0.05 | 0.9725 | 0.9774 | 0.9120 | 0.9185 | 0.8455 | 0.8529 |
| 0.2 | 0.1 | 0.9664 | 0.9737 | 0.9088 | 0.9133 | 0.8450 | 0.8498 |
| 0.2 | 0.2 | 0.9550 | 0.9609 | 0.8970 | 0.9022 | 0.8357 | 0.8397 |
| 0.2 | 0.4 | 0.9343 | 0.9345 | 0.8787 | 0.8773 | 0.8167 | 0.8158 |
| 0.4 | 0.05 | 0.9648 | 0.9687 | 0.9060 | 0.9113 | 0.8402 | 0.8455 |
| 0.4 | 0.1 | 0.9573 | 0.9584 | 0.8982 | 0.9018 | 0.8356 | 0.8387 |
| 0.4 | 0.2 | 0.9423 | 0.9391 | 0.8860 | 0.8832 | 0.8224 | 0.8206 |
| 0.4 | 0.4 | 0.8996 | 0.8734 | 0.8439 | 0.8182 | 0.7877 | 0.7605 |

Table S9: Adjacent order accuracy of SCITE for scenario 1. Branch lengths follow an exponential distribution with mean 0.2. Each cell corresponds to unique $\alpha$ and $\beta$, type of genotype and missing data percentage (Columns 3-4: data with no missing values, columns 5-6: 10% of data missing, columns 7-8: 20% of data missing).

| **FPR** $\alpha$ | **FNR** $\beta$ | **Accuracy** ternary | **Accuracy** binary | **Accuracy** 10% ternary | **Accuracy** 10% binary | **Accuracy** 20% ternary | **Accuracy** 20% binary |
|---|---|---|---|---|---|---|---|
| 0.05 | 0.05 | 0.8006 | 0.8076 | 0.5931 | 0.6329 | 0.4717 | 0.4939 |
| 0.05 | 0.1 | 0.7738 | 0.7797 | 0.5903 | 0.6232 | 0.4532 | 0.4781 |
| 0.05 | 0.2 | 0.5898 | 0.6016 | 0.4574 | 0.4831 | 0.3514 | 0.3719 |
| 0.05 | 0.4 | 0.3466 | 0.4103 | 0.2821 | 0.3349 | 0.2286 | 0.2676 |
| 0.1 | 0.05 | 0.6894 | 0.7018 | 0.5169 | 0.5531 | 0.3994 | 0.4272 |
| 0.1 | 0.1 | 0.6183 | 0.6286 | 0.4683 | 0.5089 | 0.3704 | 0.3951 |
| 0.1 | 0.2 | 0.5046 | 0.5238 | 0.3819 | 0.4234 | 0.3038 | 0.3313 |
| 0.1 | 0.4 | 0.2546 | 0.3374 | 0.2057 | 0.2829 | 0.1784 | 0.232 |
| 0.2 | 0.05 | 0.5343 | 0.5351 | 0.4069 | 0.4298 | 0.3136 | 0.3386 |
| 0.2 | 0.1 | 0.4762 | 0.4800 | 0.3594 | 0.3941 | 0.2871 | 0.3177 |
| 0.2 | 0.2 | 0.3678 | 0.3998 | 0.2828 | 0.3192 | 0.2336 | 0.2593 |
| 0.2 | 0.4 | 0.1016 | 0.2579 | 0.0881 | 0.2170 | 0.0783 | 0.1844 |
| 0.4 | 0.05 | 0.3316 | 0.3243 | 0.2572 | 0.2828 | 0.2093 | 0.2348 |
| 0.4 | 0.1 | 0.2908 | 0.2974 | 0.2211 | 0.2507 | 0.1881 | 0.2134 |
| 0.4 | 0.2 | 0.2119 | 0.2430 | 0.1664 | 0.2119 | 0.1483 | 0.178 |
| 0.4 | 0.4 | 0.0686 | 0.1669 | 0.0636 | 0.1457 | 0.0641 | 0.1333 |

Table S10: Order accuracy of SCITE for scenario 1. Branch lengths follow an exponential distribution with mean 0.2. Each cell corresponds to unique $\alpha$ and $\beta$, type of genotype and missing data percentage (Columns 3-4: data with no missing values, columns 5-6: 10% of data missing, columns 7-8: 20% of data missing).

| **FPR** $\alpha$ | **FNR** $\beta$ | **Accuracy** ternary | **Accuracy** binary | **Accuracy** 10% ternary | **Accuracy** 10% binary | **Accuracy** 20% ternary | **Accuracy** 20% binary |
|---|---|---|---|---|---|---|---|
| 0.05 | 0.05 | 0.8963 | 0.9044 | 0.8053 | 0.8361 | 0.7539 | 0.7756 |
| 0.05 | 0.1 | 0.9000 | 0.9027 | 0.8252 | 0.8475 | 0.7582 | 0.7733 |
| 0.05 | 0.2 | 0.8098 | 0.8210 | 0.7390 | 0.7628 | 0.6748 | 0.6897 |
| 0.05 | 0.4 | 0.6474 | 0.7294 | 0.5888 | 0.678 | 0.5406 | 0.6156 |
| 0.1 | 0.05 | 0.8219 | 0.8255 | 0.7495 | 0.7751 | 0.6944 | 0.7237 |
| 0.1 | 0.1 | 0.7849 | 0.7948 | 0.7153 | 0.7518 | 0.6613 | 0.6967 |
| 0.1 | 0.2 | 0.7396 | 0.7578 | 0.6769 | 0.7093 | 0.6205 | 0.6558 |
| 0.1 | 0.4 | 0.5003 | 0.6655 | 0.4628 | 0.6273 | 0.4329 | 0.5711 |
| 0.2 | 0.05 | 0.7140 | 0.7185 | 0.6590 | 0.6856 | 0.6193 | 0.6401 |
| 0.2 | 0.1 | 0.6875 | 0.6885 | 0.6283 | 0.6593 | 0.5902 | 0.6138 |
| 0.2 | 0.2 | 0.6172 | 0.6445 | 0.5633 | 0.6064 | 0.525 | 0.5738 |
| 0.2 | 0.4 | 0.2589 | 0.5653 | 0.2419 | 0.5397 | 0.2355 | 0.505 |
| 0.4 | 0.05 | 0.5825 | 0.5740 | 0.5491 | 0.5630 | 0.5187 | 0.545 |
| 0.4 | 0.1 | 0.5504 | 0.5572 | 0.5098 | 0.5393 | 0.4863 | 0.5168 |
| 0.4 | 0.2 | 0.4732 | 0.5207 | 0.4337 | 0.5076 | 0.412 | 0.4828 |
| 0.4 | 0.4 | 0.1664 | 0.4617 | 0.1709 | 0.4481 | 0.1666 | 0.4348 |

Table S11: Adjacent order accuracy of SCITE for scenario 2. Tree branch lengths follow an exponential distribution with mean 0.1. Each cell corresponds to unique $\alpha$ and $\beta$, type of genotype and missing data percentage (Columns 3-4: data with no missing values, columns 5-6: 10% of data missing, columns 7-8: 20% of data missing).

| **FPR** $\alpha$ | **FNR** $\beta$ | **Accuracy** ternary | **Accuracy** binary | **Accuracy** 10% ternary | **Accuracy** 10% binary | **Accuracy** 20% ternary | **Accuracy** 20% binary |
|---|---|---|---|---|---|---|---|
| 0.05 | 0.05 | 0.7899 | 0.8264 | 0.6115 | 0.6667 | 0.4785 | 0.5038 |
| 0.05 | 0.1 | 0.7066 | 0.7429 | 0.5608 | 0.5985 | 0.4340 | 0.4590 |
| 0.05 | 0.2 | 0.5743 | 0.6158 | 0.4591 | 0.4948 | 0.3628 | 0.3926 |
| 0.05 | 0.4 | 0.3396 | 0.4144 | 0.2809 | 0.3381 | 0.2304 | 0.2691 |
| 0.1 | 0.05 | 0.6631 | 0.7006 | 0.5130 | 0.5593 | 0.4083 | 0.4351 |
| 0.1 | 0.1 | 0.5951 | 0.6331 | 0.4684 | 0.5119 | 0.3649 | 0.4014 |
| 0.1 | 0.2 | 0.4831 | 0.5291 | 0.3876 | 0.4224 | 0.3038 | 0.3366 |
| 0.1 | 0.4 | 0.2453 | 0.3467 | 0.2125 | 0.2876 | 0.1741 | 0.2393 |
| 0.2 | 0.05 | 0.5046 | 0.5368 | 0.3989 | 0.4448 | 0.3204 | 0.3496 |
| 0.2 | 0.1 | 0.4554 | 0.4878 | 0.3618 | 0.3976 | 0.2816 | 0.3265 |
| 0.2 | 0.2 | 0.3428 | 0.3574 | 0.2876 | 0.3118 | 0.2383 | 0.2404 |
| 0.2 | 0.4 | 0.1046 | 0.2624 | 0.0921 | 0.2224 | 0.0788 | 0.1922 |
| 0.4 | 0.05 | 0.3150 | 0.3472 | 0.2528 | 0.2901 | 0.2133 | 0.2341 |
| 0.4 | 0.1 | 0.2712 | 0.3016 | 0.2282 | 0.2623 | 0.1882 | 0.2039 |
| 0.4 | 0.2 | 0.1924 | 0.245 | 0.1732 | 0.2145 | 0.1518 | 0.1794 |
| 0.4 | 0.4 | 0.0696 | 0.1668 | 0.0652 | 0.1466 | 0.0653 | 0.1319 |

Table S12: Order accuracy of SCITE for scenario 2. Tree branch lengths follow an exponential distribution with mean 0.1. Each cell corresponds to unique $\alpha$ and $\beta$, type of genotype and missing data percentage (Columns 3-4: data with no missing values, columns 5-6: 10% of data missing, columns 7-8: 20% of data missing).

| **FPR** $\alpha$ | **FNR** $\beta$ | **Accuracy** ternary | **Accuracy** binary | **Accuracy** 10% ternary | **Accuracy** 10% binary | **Accuracy** 20% ternary | **Accuracy** 20% binary |
|---|---|---|---|---|---|---|---|
| 0.05 | 0.05 | 0.8804 | 0.9148 | 0.8174 | 0.8538 | 0.7536 | 0.7804 |
| 0.05 | 0.1 | 0.8607 | 0.8859 | 0.8057 | 0.8297 | 0.7352 | 0.7545 |
| 0.05 | 0.2 | 0.7996 | 0.8298 | 0.7407 | 0.7706 | 0.6760 | 0.7016 |
| 0.05 | 0.4 | 0.6372 | 0.7386 | 0.5947 | 0.6770 | 0.5484 | 0.6220 |
| 0.1 | 0.05 | 0.8016 | 0.8315 | 0.7551 | 0.7846 | 0.7013 | 0.7249 |
| 0.1 | 0.1 | 0.7651 | 0.8034 | 0.7176 | 0.7561 | 0.6634 | 0.7025 |
| 0.1 | 0.2 | 0.7214 | 0.7618 | 0.6786 | 0.7110 | 0.6236 | 0.6554 |
| 0.1 | 0.4 | 0.4994 | 0.6789 | 0.4698 | 0.6299 | 0.4341 | 0.5820 |
| 0.2 | 0.05 | 0.6973 | 0.7226 | 0.6604 | 0.6945 | 0.6191 | 0.6526 |
| 0.2 | 0.1 | 0.6684 | 0.6981 | 0.6275 | 0.6640 | 0.5874 | 0.6254 |
| 0.2 | 0.2 | 0.5901 | 0.6189 | 0.5605 | 0.5914 | 0.5274 | 0.5315 |
| 0.2 | 0.4 | 0.2606 | 0.5811 | 0.2507 | 0.5428 | 0.2434 | 0.5108 |
| 0.4 | 0.05 | 0.5686 | 0.5971 | 0.5469 | 0.5797 | 0.5217 | 0.5423 |
| 0.4 | 0.1 | 0.5345 | 0.5652 | 0.5062 | 0.5480 | 0.4851 | 0.5206 |
| 0.4 | 0.2 | 0.4520 | 0.5272 | 0.4333 | 0.5147 | 0.4234 | 0.4840 |
| 0.4 | 0.4 | 0.1753 | 0.4655 | 0.1727 | 0.4482 | 0.1719 | 0.4303 |

Table S13: Adjacent order accuracy of SiFit for scenario 1. Tree branch lengths follow an exponential distribution with mean 0.2. Each cell corresponds to unique $\alpha$ and $\beta$, type of genotype and missing data percentage (Columns 3-4: data with no missing values, columns 5-6: 10% of data missing, columns 7-8: 20% of data missing).

| **FPR** $\alpha$ | **FNR** $\beta$ | **Accuracy** ternary | **Accuracy** binary | **Accuracy** 10% ternary | **Accuracy** 10% binary | **Accuracy** 20% ternary | **Accuracy** 20% binary |
|---|---|---|---|---|---|---|---|
| 0.05 | 0.05 | NA | 0.3150 | NA | 0.2278 | NA | 0.1482 |
| 0.05 | 0.1 | NA | 0.3216 | NA | 0.2298 | NA | 0.1484 |
| 0.05 | 0.2 | NA | 0.2039 | NA | 0.1449 | NA | 0.0963 |
| 0.05 | 0.4 | NA | 0.1202 | NA | 0.0864 | NA | 0.0582 |
| 0.1 | 0.05 | NA | 0.3184 | NA | 0.2205 | NA | 0.1381 |
| 0.1 | 0.1 | NA | 0.2669 | NA | 0.1854 | NA | 0.1191 |
| 0.1 | 0.2 | NA | 0.2012 | NA | 0.1411 | NA | 0.0921 |
| 0.1 | 0.4 | NA | 0.0998 | NA | 0.0725 | NA | 0.0479 |
| 0.2 | 0.05 | NA | 0.1820 | NA | 0.1268 | NA | 0.0789 |
| 0.2 | 0.1 | NA | 0.1398 | NA | 0.1020 | NA | 0.0681 |
| 0.2 | 0.2 | NA | 0.0885 | NA | 0.0618 | NA | 0.0423 |
| 0.2 | 0.4 | NA | 0.0371 | NA | 0.0276 | NA | 0.0199 |
| 0.4 | 0.05 | NA | 0.0216 | NA | 0.0120 | NA | 0.0082 |
| 0.4 | 0.1 | NA | 0.0148 | NA | 0.0089 | NA | 0.0043 |
| 0.4 | 0.2 | NA | 0.0066 | NA | 0.0050 | NA | 0.0031 |
| 0.4 | 0.4 | NA | 0.0069 | NA | 0.0062 | NA | 0.0045 |

Table S14: Order accuracy of SiFit for scenario 1. Tree branch lengths follow an exponential distribution with mean 0.2. Each cell corresponds to unique $\alpha$ and $\beta$, type of genotype and missing data percentage (Columns 3-4: data with no missing values, columns 5-6: 10% of data missing, columns 7-8: 20% of data missing).

| **FPR** $\alpha$ | **FNR** $\beta$ | **Accuracy** ternary | **Accuracy** binary | **Accuracy** 10% ternary | **Accuracy** 10% binary | **Accuracy** 20% ternary | **Accuracy** 20% binary |
|---|---|---|---|---|---|---|---|
| 0.05 | 0.05 | NA | 0.2587 | NA | 0.1958 | NA | 0.1428 |
| 0.05 | 0.1 | NA | 0.2604 | NA | 0.1945 | NA | 0.1399 |
| 0.05 | 0.2 | NA | 0.1852 | NA | 0.1391 | NA | 0.1027 |
| 0.05 | 0.4 | NA | 0.1179 | NA | 0.0929 | NA | 0.0687 |
| 0.1 | 0.05 | NA | 0.2601 | NA | 0.1898 | NA | 0.1325 |
| 0.1 | 0.1 | NA | 0.2264 | NA | 0.1639 | NA | 0.1171 |
| 0.1 | 0.2 | NA | 0.1794 | NA | 0.1348 | NA | 0.0958 |
| 0.1 | 0.4 | NA | 0.1061 | NA | 0.0800 | NA | 0.0589 |
| 0.2 | 0.05 | NA | 0.1482 | NA | 0.1139 | NA | 0.0837 |
| 0.2 | 0.1 | NA | 0.1205 | NA | 0.0965 | NA | 0.0723 |
| 0.2 | 0.2 | NA | 0.0825 | NA | 0.0659 | NA | 0.0517 |
| 0.2 | 0.4 | NA | 0.0432 | NA | 0.0347 | NA | 0.0258 |
| 0.4 | 0.05 | NA | 0.0233 | NA | 0.0153 | NA | 0.0112 |
| 0.4 | 0.1 | NA | 0.0190 | NA | 0.0130 | NA | 0.0077 |
| 0.4 | 0.2 | NA | 0.0112 | NA | 0.0082 | NA | 0.0058 |
| 0.4 | 0.4 | NA | 0.0070 | NA | 0.0069 | NA | 0.0062 |

Table S15: Adjacent order accuracy of SiFit for scenario 2. Tree branch lengths follow an exponential distribution with mean 0.1. Each cell corresponds to unique $\alpha$ and $\beta$, type of genotype and missing data percentage (Columns 3-4: data with no missing values, columns 5-6: 10% of data missing, columns 7-8: 20% of data missing).

| **FPR** $\alpha$ | **FNR** $\beta$ | **Accuracy** ternary | **Accuracy** binary | **Accuracy** 10% ternary | **Accuracy** 10% binary | **Accuracy** 20% ternary | **Accuracy** 20% binary |
|---|---|---|---|---|---|---|---|
| 0.05 | 0.05 | NA | 0.3255 | NA | 0.2368 | NA | 0.1574 |
| 0.05 | 0.1 | NA | 0.2749 | NA | 0.1996 | NA | 0.1304 |
| 0.05 | 0.2 | NA | 0.2068 | NA | 0.1511 | NA | 0.0979 |
| 0.05 | 0.4 | NA | 0.1148 | NA | 0.0861 | NA | 0.0550 |
| 0.1 | 0.05 | NA | 0.3171 | NA | 0.2202 | NA | 0.1408 |
| 0.1 | 0.1 | NA | 0.2654 | NA | 0.1858 | NA | 0.1168 |
| 0.1 | 0.2 | NA | 0.1977 | NA | 0.1354 | NA | 0.0911 |
| 0.1 | 0.4 | NA | 0.0991 | NA | 0.0677 | NA | 0.0478 |
| 0.2 | 0.05 | NA | 0.1833 | NA | 0.1284 | NA | 0.0796 |
| 0.2 | 0.1 | NA | 0.1367 | NA | 0.1004 | NA | 0.0645 |
| 0.2 | 0.2 | NA | 0.0889 | NA | 0.0632 | NA | 0.0467 |
| 0.2 | 0.4 | NA | 0.0374 | NA | 0.0278 | NA | 0.0169 |
| 0.4 | 0.05 | NA | 0.0201 | NA | 0.0121 | NA | 0.0075 |
| 0.4 | 0.1 | NA | 0.0138 | NA | 0.0081 | NA | 0.0053 |
| 0.4 | 0.2 | NA | 0.0057 | NA | 0.0046 | NA | 0.0028 |
| 0.4 | 0.4 | NA | 0.0066 | NA | 0.0053 | NA | 0.0047 |

Table S16: Order accuracy of SiFit for scenario 2. Tree branch lengths follow an exponential distribution with mean 0.1. Each cell corresponds to unique $\alpha$ and $\beta$, type of genotype and missing data percentage (Columns 3-4: data with no missing values, columns 5-6: 10% of data missing, columns 7-8: 20% of data missing).

| **FPR** $\alpha$ | **FNR** $\beta$ | **Accuracy** ternary | **Accuracy** binary | **Accuracy** 10% ternary | **Accuracy** 10% binary | **Accuracy** 20% ternary | **Accuracy** 20% binary |
|---|---|---|---|---|---|---|---|
| 0.05 | 0.05 | NA | 0.2636 | NA | 0.2031 | NA | 0.1501 |
| 0.05 | 0.1 | NA | 0.2299 | NA | 0.1755 | NA | 0.1287 |
| 0.05 | 0.2 | NA | 0.1845 | NA | 0.1427 | NA | 0.1056 |
| 0.05 | 0.4 | NA | 0.1205 | NA | 0.0933 | NA | 0.0677 |
| 0.1 | 0.05 | NA | 0.2593 | NA | 0.1866 | NA | 0.1314 |
| 0.1 | 0.1 | NA | 0.2255 | NA | 0.1645 | NA | 0.118 |
| 0.1 | 0.2 | NA | 0.1795 | NA | 0.13 | NA | 0.0956 |
| 0.1 | 0.4 | NA | 0.1018 | NA | 0.0785 | NA | 0.0583 |
| 0.2 | 0.05 | NA | 0.1497 | NA | 0.114 | NA | 0.0819 |
| 0.2 | 0.1 | NA | 0.117 | NA | 0.0932 | NA | 0.0696 |
| 0.2 | 0.2 | NA | 0.081 | NA | 0.0659 | NA | 0.0512 |
| 0.2 | 0.4 | NA | 0.0446 | NA | 0.0344 | NA | 0.025 |
| 0.4 | 0.05 | NA | 0.0238 | NA | 0.016 | NA | 0.0097 |
| 0.4 | 0.1 | NA | 0.0183 | NA | 0.012 | NA | 0.0084 |
| 0.4 | 0.2 | NA | 0.0102 | NA | 0.0075 | NA | 0.0052 |
| 0.4 | 0.4 | NA | 0.0073 | NA | 0.0065 | NA | 0.0056 |
